# High-throughput mapping of single-neuron projection and molecular features by retrograde barcoded labelling

**DOI:** 10.1101/2021.05.16.444258

**Authors:** Peibo Xu, Jian Peng, Tingli Yuan, Zhaoqin Chen, Hui He, Ziyan Wu, Ting Li, Xiaodong Li, Luyue Wang, Wu Wei, Chengyu T. Li, Zhen-Ge Luo, Yuejun Chen

## Abstract

Deciphering patterns of connectivity between neurons in the brain is a critical step toward understanding brain function. Imaging-based neuroanatomical tracing identifies area-to-area or sparse neuron-to-neuron connectivity patterns, but with limited throughput. Barcode-based connectomics maps large numbers of single-neuron projections, but remains a challenge for jointly analyzing single-cell transcriptomics. Here, we established a rAAV2-retro barcode-based multiplexed tracing method that simultaneously characterizes the projectome and transcriptome at the single neuron level. We uncovered dedicated and collateral projection patterns of ventromedial prefrontal cortex (vmPFC) neurons to five downstream targets and found that projection-defined vmPFC neurons are molecularly heterogeneous. We identified transcriptional signatures of projection-specific vmPFC neurons, and verified *Pou3f1* as a marker gene of neurons extending collateral branches to dorsomedial striatum and lateral hypothalamus. In summary, we have developed a new multiplexed technique whose paired connectome and gene expression data can help reveal organizational principles that form neural circuits and process information.

## Introduction

Wiring diagrams of a brain can be divided into three levels: 1) the macroscale connectome that describes inter-areal connections, 2) the mesoscale connectome that describes connections between cells, and 3) the microscale connectome that describes connections at the synaptic level (Zeng, 2018). Studying circuit architecture at the level of the mesoscale connectome describes how information flows between brain regions (Oh et al., 2014). Traditionally, neuroanatomical tracers are used to characterize regional connectivity matrices (Cowan, 1998). To obtain cell-type-specific connectivity, one can use recombinant virus-based tracer in transgenic model organisms or more precisely trace a specific component of a neural circuit using viral-genetic tracing tools to dissect the input-output organization (Ghosh et al., 2011; Nassi et al., 2015; Schwarz et al., 2015). However, these methods are highly reliant on complex recombinant virus design and genetically modified model organism, and often are not at a single-neuron resolution.

Achieving a single-neuron resolution connectome will undoubtedly transform our understanding of functional brain units. Previous studies have used high-throughput fluorescence microscopy to reconstruct the detailed morphology of single neurons (Gong et al., 2016; Rompani et al., 2017). In addition, barcode-based circuit mapping techniques can now harness the power of next-generation sequencing to study single-neuron projection patterns in a high-throughput manner (Han et al., 2018; Kebschull et al., 2016; Zador et al., 2012). These advances have revolutionized our understanding of neuronal connection complexity, but we still lack a comprehensive connectome by considering the roles of transcriptome-defined neuronal subtypes and their effect on input/output transformations (Fornito et al., 2019; Klingler et al., 2019; Sorensen et al., 2015). A recently developed method assessed the relationship between projection pattern and gene expression for single intratelencephalic neurons. However, the number of genes that can be interrogated is limited as it utilizes highly multiplexed single-molecule fluorescence in situ hybridization (smFISH) (Chen et al., 2019).

Medial prefrontal cortex (mPFC) is an intricate brain region involved in higher order cognitive functions, information processing (e.g., memory and emotions) and driving goal-directed actions (Le Merre et al., 2021). For example, mPFC neurons projecting to the nucleus accumbens encoding punishment-related internal states were located in more superficial layer 5a, and mPFC neurons projecting to the ventral tegmental area encoding aversive learning were located in deeper layer 5b (Kim et al., 2017; Wu et al., 2021). Though previous studies have extensively investigated the anatomical and functional diversities of mPFC, the relationship between anatomical and molecular features of mPFC neurons remains elusive. Do mPFC neurons projecting to different downstream brain regions differ in their transcriptomes? Are these projection-defined mPFC neurons homogeneous or composed of different neuron subtypes? The answer to these questions may be further complicated by the finding that mPFC neurons can send collateral axons to multiple brain regions (Cornwall and Phillipson, 1988). So, what are the principles of target selection or target combination for these collateral projection mPFC neurons? What are the cell type and molecular features of these “broadcasting” neurons?

To address these challenges, we designed a multiplexed tracing method capable of characterizing single-neuron transcriptome and projectome at the same time, which we called MERGE-seq (Multiplexed projEction neuRons retroGrade barcodE). We used MERGE-seq to interrogate the projectome and the corresponding transcriptome of ventral mPFC neurons. We injected five rAAV2-retro viruses with distinct barcodes into the five known downstream targets of ventromedial prefrontal cortex (vmPFC), including agranular insular cortex (AI), dorsomedial striatum (DMS), basal amygdala (BLA), mediodorsal thalamic nucleus (MD) and lateral hypothalamus (LH), in the same mouse brain such that each target region received a unique barcoded rAAV2-retro. We found that vmPFC neurons projecting to each downstream target are heterogeneous, which are composed of transcriptionally different subtypes of neurons. About 74% of barcoded vmPFC neurons projected to one of these five targets (dedicated projection), and ∼26% of barcoded vmPFC neurons sent collateral projections to multiple brain regions, most of which are dual-target projection neurons (bifurcated projection). We further uncovered the cell type compositions and layer distributions of these dedicated and bifurcated projection vmPFC neurons, and revealed their molecular signatures. By combination of RNA fluorescence in situ hybridization (FISH) and dual-color retrograde AAV labelling, we verified the bifurcated projection of vmPFC neurons, and demonstrated that *Pou3f1*^+^ neurons in layer 5 can collaterally project to DMS and LH. Finally, we implemented a machine learning-based methodology and uncovered specific gene clusters for predicting certain projection patterns. As MERGE-seq bridges the gap between single-neuron projectome and transcriptome data, it can uncover new molecular properties of anatomical neural circuits.

## Results

### MERGE-seq characterizes single neuron transcriptome and projectome simultaneously

In order to use the 10X Genomics scRNA-seq system to analyze transcripts from cells infected with rAAV2-retro virus, we modified the viral vector by adding a 15 bp barcode index and polyadenylation signal sequences to the 3’ end of the EGFP sequences, which was driven by a short CAG promoter (***Figure 1A, B***, see **Methods)**. Then, five rAAV2-retro viruses with different barcodes were individually injected into five brain regions of the same mouse, including AI, DMS, BLA, LH, and MD. These brain areas are the known downstream brain regions of vmPFC (Hunnicutt et al., 2016; Hurley et al., 1991; Reppucci and Petrovich, 2016; Vertes, 2004; Zhu et al., 2020). A period of six weeks was set to allow efficient retrograde labelling of vmPFC neurons by these barcoded viruses. These mice were then sacrificed and the vmPFC (specifically the prelimbic area (PrL) and the infralimbic area (IL)) was carefully dissected for scRNA-seq analysis (***Figure 1A***). Single-cell transcriptional libraries were obtained using 10X Genomics library preparation protocols, and virus barcode expression libraries were obtained using user-defined primers, which could enrich cDNA fragments composed of barcode index, unique molecular identifiers (UMIs), and the cell barcode (***Figure 1B***). We recovered 1791 EGFP-positive cells undergoing fluorescence-activated cell sorting (FACS) from three mice and 19,470 single cells without sorting from the other three mice, a total of 21,261 cells. Transcriptional profiling of all cells revealed major cell types including excitatory neurons (*Slc17a7*^+^), microglia (*C1qa*^+^), endothelial cells (*Itm2a*^+^, Endo), oligodendrocyte progenitor cells (*Olig2*^+^*Mog*^-^, OPCs), oligodendrocyte (*Olig2*^+^*Mog*^+^, Oligo), inhibitory neurons (*Gad1*^+^), astrocyte (*Aldh1l1*^+^, Astro) and activated microglia (*C1qa*^+^*Pf4*^+^, Act. Microglia) as previously reported (Bhattacherjee et al., 2019) (***Figure 1C-E***). Barcoded cells below refer to a collection of barcoded cells from unsorted group and FAC-sorted group.

**Figure 1.**
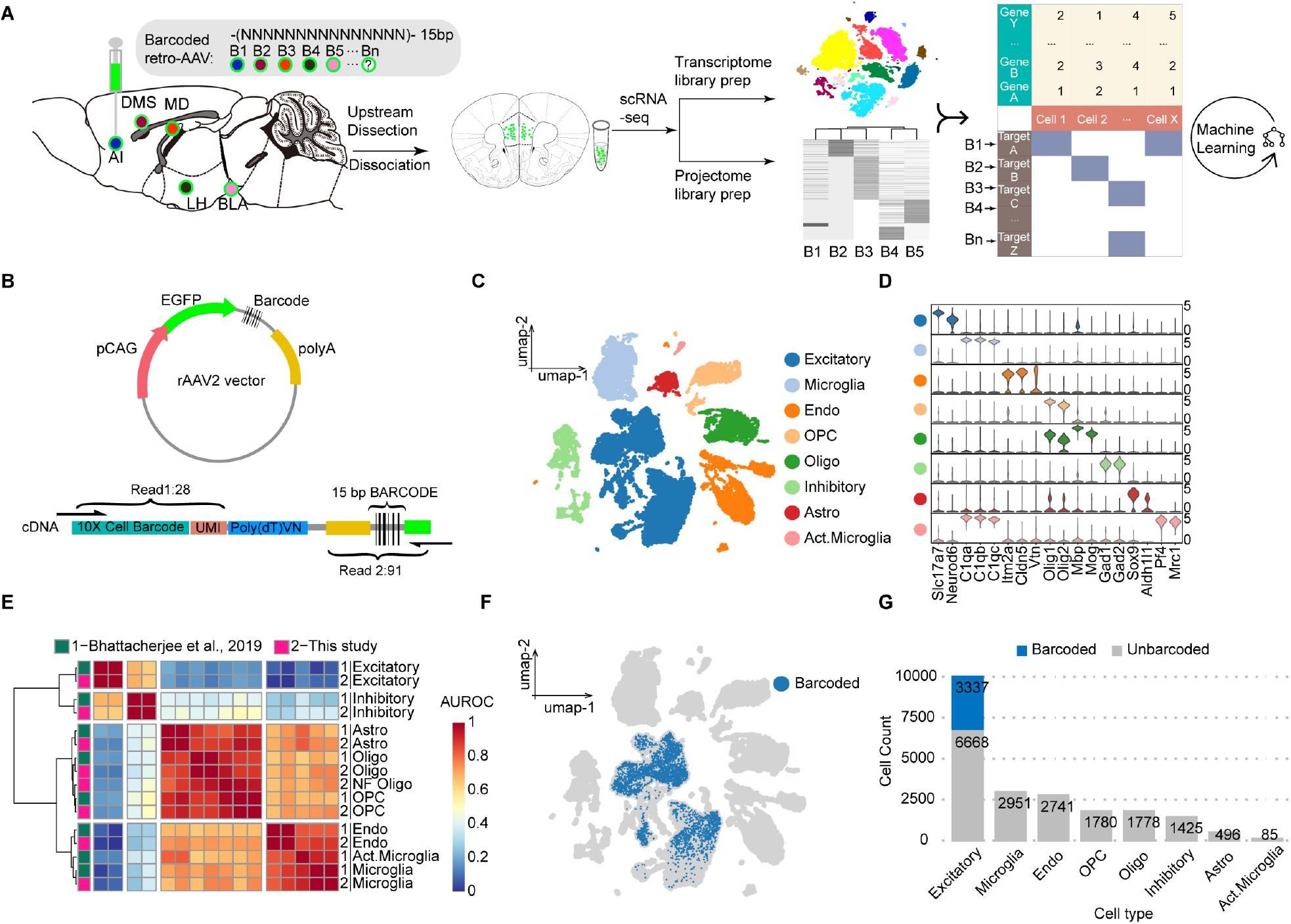
MERGE-seq characterized single-neuron transcriptomes and projectomes simultaneously. (**A**) Schematic diagram of the experimental workflow. (**B**) rAAV2 plasmid vector design, and schematic of designed primers to recover cell barcode and UMI in read 1, and 3’ tail of EGFP and virus barcode in read 2. According to the recommendation of 10X Genomics, a faithful mapping should cover 28 bp for read 1 and 91 bp for read 2. In our design, 150 bp pair-end sequencing can sufficiently meet the need to recover cell barcode, UMI and virus barcode. (**C**) Umap embedding of transcriptional clustering results for all vmPFC cells. (**D**) Stacked violin plots showing the expression of markers for each cluster. (**E**) Heatmap showing correlation between cell types recovered in this study and the dataset from Bhattacherjee et al. (Bhattacherjee et al., 2019) (**F**) Umap embedding of all determined barcoded cells labeled in blue. (**G**) Bar plot showing frequency of barcoded (blue) and unbarcoded (grey) cells in all recovered cell types.

First, we validated that each target region was labeled and effectively covered by the rAAV2-retro-EGFP (***Figure 1-figure supplement 1A***). Next, we showed that there were sufficient sequence differences to distinguish one barcode to others and sufficient sequence difference to identify the right barcode among 5 references during sequencing (***Figure 1-figure supplement 1B, C***). Since each downstream brain region of vmPFC received a unique and predetermined barcoded virus, each virus barcode identified in a vmPFC neuron represents the specific corresponding downstream brain region that the neuron projects to. To confidently determine the virus barcode identities of recovered cells, we hypothesized that non-neuron cells would not be transduced by rAAV2-retro and calculated the empirical cumulative distribution of UMI counts of non-neuron cells for each virus barcode. We then set a stringent cumulative density threshold of 0.999, where corresponding UMI counts were determined as filtering threshold (see **Methods, *Figure 1-figure supplement 1D-I***). Across all detected cell types, barcoded cells were primarily excitatory neurons rather than inhibitory neurons or non-neuronal cell types (3337 validly barcoded in 10005 excitatory neurons, and 0 validly barcoded in other cell types, ***Figure 1F, G***). This is consistent with the finding that mPFC projection neurons are excitatory (Gabbott et al., 2005). Based on this stringent threshold, we found that the percentage of barcoded cells in FAC-sorted or unsorted groups is 54% and 12%, respectively (***Figure 1-figure supplement 1J***). Together, these results demonstrate that MERGE-seq can record single neuron transcriptome and projectome simultaneously.

### MERGE-seq reveals transcriptomic heterogeneity and cell type composition of vmPFC neurons projecting to different targets

Previous studies have shown that vmPFC neurons project to multiple brain regions including AI, DMS, BLA, LH, and MD, however, the cell type composition of these projection neurons remains largely unknown (Le Merre et al., 2021). Combining with single neuron transcriptome, we explored the transcriptome and subtype composition of vmPFC neurons projecting to different downstream brain regions. We first performed clustering analysis on excitatory projection neurons expressing *Slc17a7* (also known as vesicular glutamate transporter, *Vglut1*) and filtered out stressed 637 neurons (***Figure 2-figure supplement 1A***). We generated 7 excitatory neuron clusters, which were annotated based on typical markers of cortical layers (Bhattacherjee et al., 2019; Sorensen et al., 2015) (layer 2/3, *Cux2*; layer 5, *Etv1*; layer 6, *Sulf1*) and differentially expressed genes (DEGs) (***Supplementary file 1***). These neuron clusters include L2/3-Calb1 (3.8%), L2/3-Rorb (7.4%), L5-Bcl6 (4.1%), L5-Htr2c (3.4%), L5-S100b (15%), L6-Npy (11.9%) and L6-Syt6 (54.4%) (***Figure 2A***). The layer and subtype marker genes of these clusters were confirmed to be expressed in corresponding layers in the vmPFC, as revealed by in situ hybridization results of the Allen Mouse Brain Atlas (***Figure 2A, Figure2-figure supplement 1A-C***). Of note, we captured more layer 6 neurons than superficial layer neurons (11.9% L6-Npy and 54.4% L6-Syt6, ***Figure 2B***), which is different from a previous report (Bhattacherjee et al., 2019). We speculate that different dissociation protocols may cause biased neuron capture.

**Figure 2.**
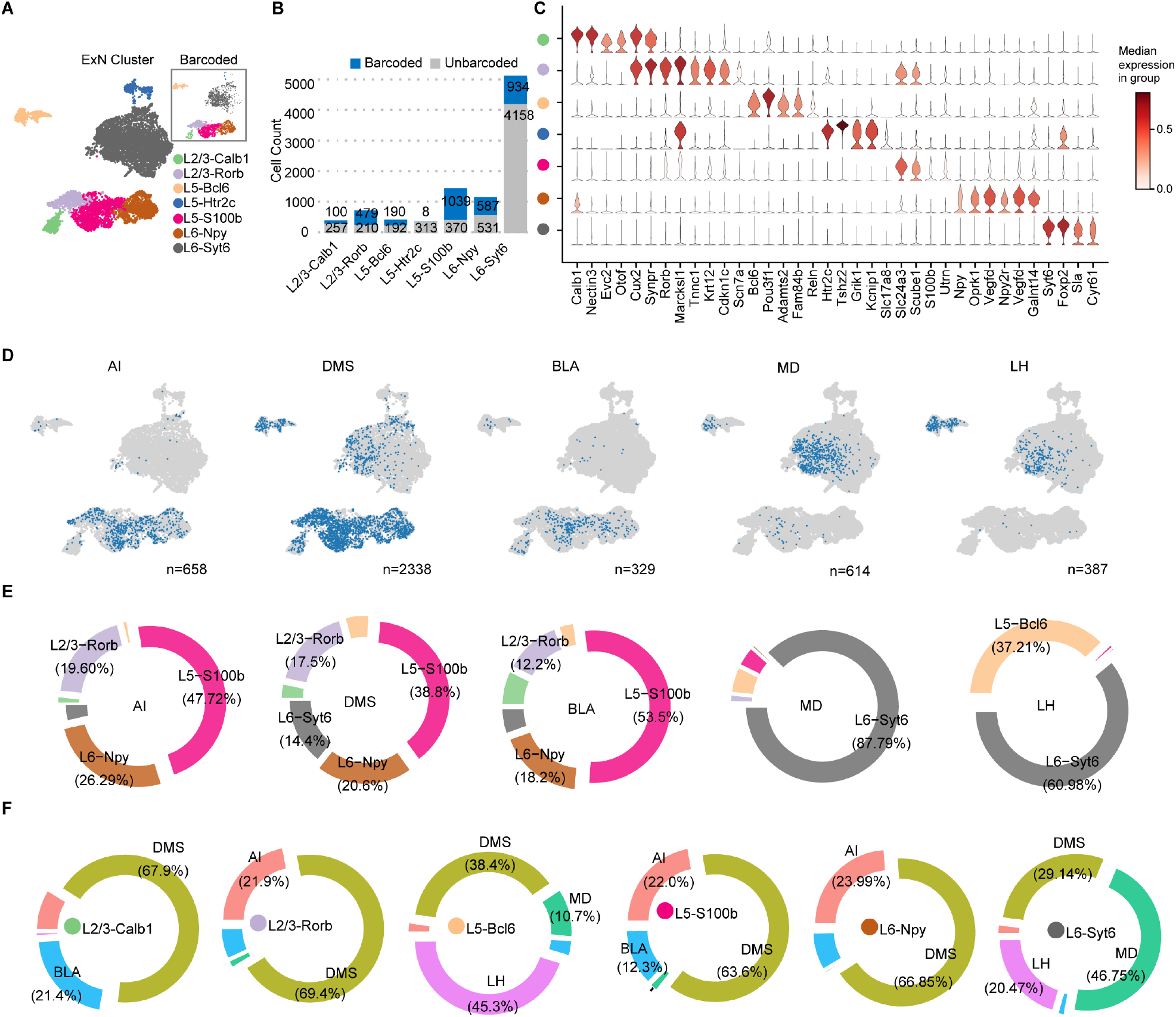
MERGE-seq unravels transcriptomic heterogeneity of projection target-defined vmPFC neurons. (**A**) Umap embedding of excitatory neuron subtype annotation. (**B**) Bar plot showing frequency of barcoded (blue) and unbarcoded (grey) neurons in distinct neuron subtypes. (**C**) Stacked violin plot showing the expression of markers for each neuronal subtype. (**D**) Umap embeddings of barcoded (blue) neurons projecting to each target. (**E**) Donut plot describing the distribution of neuronal subtypes (donut) for barcoded neurons associated with each projection target. Threshold to show label as a ratio of total was set to 0.1. Neuronal subtype color codes are the same as in (**A**). (**F**) Donut plot describing the distribution of projection targets (donut) for barcoded neurons associated with each neuronal type. Threshold to show label as a ratio of total was set to 0.1.

Cells that were retrogradely barcoded spanned all layers of the vmPFC (layer 2/3, 5, and 6) and included all 7 neuronal subtypes (***Figure 2A-C***). These subtypes were transcriptionally similar to those previously reported (***Figure2-figure supplement 1D, E***), suggesting that multiple viral infections will not significantly affect the transcriptional state of these retrogradely labeled vmPFC neurons (Bhattacherjee et al., 2019). For the L5-Htr2c subtype, only 8 neurons were validly barcoded, indicating that these neurons usually do not project to any target we chose (***Figure 2B***). Neurons projecting to DMS were abundant (n = 2338), whereas neurons projecting to BLA were rare (n = 329) (***Figure 2D***). These results are consistent with data acquired via conventional fluorescence-based retrograde tracing in the prefrontal cortex of rats (Gabbott et al., 2005).

We next calculated the subtype composition of vmPFC neurons projecting to each downstream brain regions. Interestingly, we found that these target specific projection neurons were transcriptionally heterogeneous, which were composed of different neuronal subtypes (***Figure 2D, E***). Neurons projecting to LH or MD were mainly L6-Syt6 subtype (***Figure 2E***), whereas neurons projecting to AI, DMS, or BLA were mainly composed of L5-S100b, and to a lesser extent L6-Npy and L2/3-Rorb subtypes (***Figure 2E***). It is worth noting that the cellular composition of target specific projection neurons from FAC-sorted or unsorted groups is comparable (***Figure2-figure supplement 1F***).

As the layer distribution of each neuron subtype can be inferred by their layer specific marker gene expression, these results also implied the layer distribution of neurons projecting to each target (***Figure 2E, Figure2-figure supplement 1B***). By calculating the projection properties of each vmPFC neuron subtypes, we found that each transcriptome-defined neuron subtype can project to specific but multiple targets. For instance, L5-S100b, L6-Npy and L2/3-Rorb mainly projected to AI, DMS and BLA, while L6-Syt6 mainly projected to MD and LH (***Figure 2F***). Interestingly, we also found that different neuron subtypes localized in the same layer could project to distinct targets. For instance, L6-Npy neurons projecting to AI, DMS and BLA, while L6-Syt6 neurons projecting to MD, DMS and LH (***Figure 2F***). Similar phenotypes were observed for L5-S100b and L5-Bcl6 subtypes (***Figure 2F***), suggesting transcriptomic and projection/function diversities in the spatially close neurons within the same cortical layer.

Together, by MERGE-seq analysis, we have revealed the heterogeneity and cellular composition of vmPFC neurons projecting to different target. Our results demonstrate that vmPFC neurons projecting to a certain target are composed of different transcriptome-defined neuron subtypes, and individual transcriptome-defined subtypes of vmPFC neuron project to multiple targets.

### MERGE-seq reveals dedicated and collateral projection patterns of vmPFC neuron at single cell level

Interestingly, we found that a portion of barcoded vmPFC neurons had more than one type of barcode, suggesting collateral projection of these neurons. We therefore analyzed the projection pattern of each barcoded vmPFC neuron by calculating the number of valid barcode types (see **Methods**). We defined the dedicated projection neuron as a neuron containing only one type of barcode, the collateral projection neuron as a neuron containing more than one type of barcode. We found 74.35% of 3183 viral-barcoded neurons belonged to dedicated projection and the remaining belonged to collateral projection. 22.00% had dual targets (bifurcated projection), 3.36% had triple targets, and 0.30%, if any, projected to more than three targets (***Figure 3A***). By calculating the conditional probability that the same neuron projects to two targets (see **Methods**), we found that vmPFC neurons projecting to AI or BLA were more likely to have collateral projection to DMS (***Figure 3B***). We also observed a relatively high conditional probability of collateral projection between MD and LH, or DMS and LH, or DMS and MD (***Figure 3B***), suggesting bifurcated projections to these paired targets for single vmPFC neuron. We further compared our experimental results to a null model assuming the independent projection probability of each neuron to different targets, simply as a binomial distribution of projection (Gergues et al., 2020; Han et al., 2018). We found that 2 projection patterns were over-represented and 8 were under-represented after adjusting for multiple comparisons (Bonferroni correction, ***Supplementary file 2, Figure 3C, D***). All projection patterns comprised of more than three targets were not over- or under-represented (***Figure 3C, D***). These 10 over- or under-represented projection patterns were ∼32% of the total (10/31), suggesting that vmPFC neurons project to a subset of these five targets in a non-random manner.

**Figure 3.**
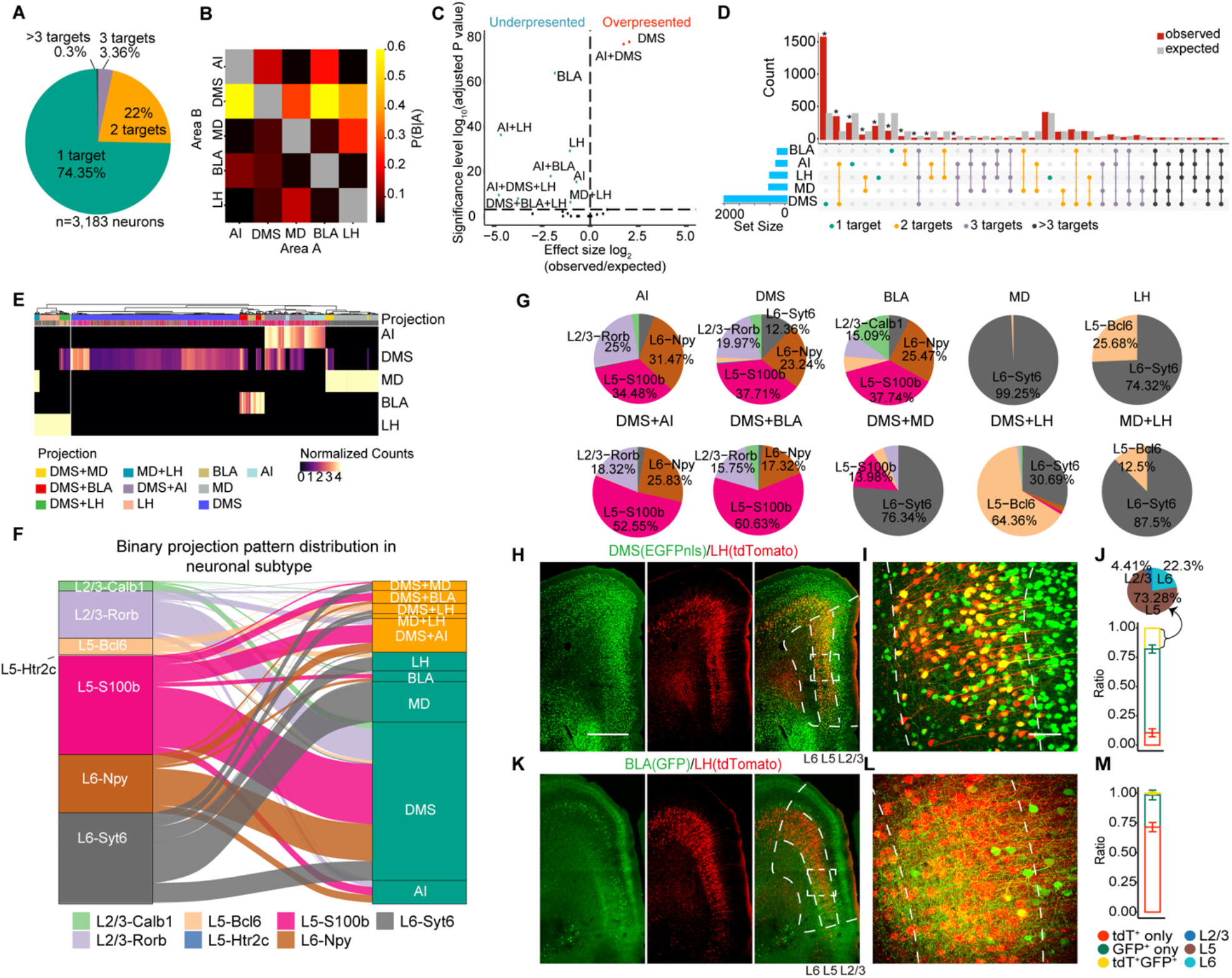
MERGE-seq reveals hidden projection diversity within the vmPFC. (**A**) Pie chart indicating the number of projection targets for barcoded vmPFC neurons recovered by MERGE-seq. (**B**) Heatmap showing the probability that a neuron projecting to area A also projects to area B. (**C**) Volcano plot of adjusted statistical significance level and effect size of over- and under-represented projection patterns (two-sided Binomial test, *p-value ≤0.001, adjusted P values and effect sizes are provided in table S2). (**D**) Upset plot showing the intersection of five projection targets. Horizontal blue bars represent the frequency of barcoded cells for each projection target. Vertical red and grey bars represent observed and expected neurons of each projection pattern, respectively. Comparisons were performed between observed and expected neurons using two-sided binomial tests. P values were adjusted using the Bonferroni method. (**E**) Heatmap showing normalized projection strength. Rows represent the projection targets and columns represent the cells labeled by the top 10 binary projection patterns or labeled by transcriptional neuron subtypes. (**F**) Alluvial plot showing the 10 most frequent projection patterns distribution into neuronal subtypes. (**G**) Pie charts describing the projection patterns from (**F**) partitioned by neuronal subtype. Only ratios greater than 10% are shown. (**H-M**) Immunostaining of dual-color traced retrograde labeled neurons of selected targets (DMS+LH and BLA+LH) (**H, K**). Dotted line depicts layers 2/3, 5, and 6 of the vmPFC. Scale bars, 500 μm. (**I, L**) Enlarged view of the dotted box in (**H, K**). Scale bars, 100 μm. (**J, M**) Histogram shows quantitative data for single-(red, green) and double- (yellow) labeled neurons as mean percentages of total rAAV2-retro labeled neurons (n = 3 mice). Data are presented as mean ± SD. Pie chart showing layer distribution of double (yellow) labeled neurons.

We then focused our analysis on the 5 dedicated projections (DMS, AI, MD, LH, and BLA) and most frequent five bifurcated projections (DMS+AI, MD+LH, DMS+MD, DMS+BLA, and DMS+LH). We conducted a principal component analysis (PCA) of the projection matrix and mapped binary projection labels on PC embeddings. Results from binary projection clustering aligned well with clusters at PC1- and PC2-defined embeddings, except for a scattered distribution of the DMS/BLA bifurcated group (***Figure3-figure supplement 1A***). We further clustered cells according to projection strength (defined as normalized virus barcode UMI counts) (***Figure 3E***). We found that cells had bifurcated projections to DMS+MD, or MD+LH, or DMS+AI, or DMS+LH (***Figure 3E***), a pattern very similar to that we observed in binary projection model, indicating that projection strength-based clustering is comparable to binary projection pattern model (***Figure 3B***).

We next explored the cell type composition of the top 10 dedicated or bifurcated projection neurons. We mapped transcriptomic clusters to projection patterns (***Figure 3F***). We found that most dedicated or bifurcated projection neurons are transcriptionally diverse, consisting of at least three neuron subtypes (***Figure 3G***), whereas MD-projecting, LH-projecting, and MD+LH-projecting cells are much more homogenous, mainly composed of L6-Syt6 neurons (***Figure 3G***).

To validate the bifurcated projection patterns inferred from the digital projectome, we injected retrograde AAV2 encoding different fluorescent proteins (EGFP or tdTomato) into different combinations of projection targets (dual-color rAAV2-retro labelling assay), and analyzed the projection patterns by immunohistochemistry. We observed 17.8% ± 0.11% dual-color labeled neurons of all virus labeled neurons in DMS+LH group, suggesting bifurcated projection of vmPFC neurons to these two targets (***Figure 3H-J***). Furthermore, 73.28% ± 7.60% of total dual-color labeled neurons were located in layer 5 (***Figure 3H-J***). These results are consistent with our digital projectome map revealed by MERGE-seq, showing that vmPFC neurons have bifurcated projections to DMS and LH (***Figure 3B-F***), and DMS+LH-projecting neurons are mainly L5-Bcl6 subtype, which were located in layer 5 (***Figure 3G***). Other bifurcated projection patterns inferred by MERGE-seq was also verified by our dual-color retro-AAV labelling assay. These patterns included DMS+AI (23.1% ± 2.03% of all dual-color neurons) and DMS+BLA (6.59% ± 1.55%) (***Figure1-figure supplement 1B-D***). In contrast, we only observed 1.66% ± 0.92% of dual-color labeled neurons in BLA+LH group (***Figure 3K-M***). This result is consistent with our MERGE-seq analysis, in which BLA+LH was not inferred as bifurcated projection targets (***Figure 3B-E***), further supporting the accuracy of the digital projectome based on MERGE-seq analysis. This result also supports the view that vmPFC neuron projects (including collaterally project) to downstream targets in a non-random manner (***Figure 3C, D***).

Altogether, by MERGE-seq analysis, we have revealed the projection diversities of vmPFC neurons at single neuron level. These results demonstrate that single vmPFC neuron has dedicated or collateral projections, and the projection pattern of these neurons are specific, but not in a random manner. Furthermore, projection-defined (dedicated or bifurcated) neurons have specific cell type composition and layer distributions. It is worth noting that as a proof of concept, we only acquired the vmPFC projectome from five downstream targets. Definitions to dedicated or collateral projections are thus limited to these five targets and some collateral projections may be underestimated.

### Transcriptional profiling of projection target-specific vmPFC neurons

Next, we sought to determine the molecular features of neurons projecting to different downstream targets. We calculated differentially expressed genes (DEGs) for each target specific projection neurons (***Figure 4A, B, Figure4-figure supplement 1***). We found that some of projection-specific DEGs are marker genes of typical neuronal types. For example, *Syt6, Foxp2, Sla* and *Cyr61* are both MD-projecting DEGs and marker genes of L6-Syt6 neurons; *Marcksl1, Rorb* and *Cux2* are both DMS-projecting DEGs and marker genes of layer 2/3 neurons (neuronal subtypes L2/3-Calb1 and L2/3-Rorb) (***Figure 2C, Figure 4A, B, Figure4-figure supplement 1***).

**Figure 4.**
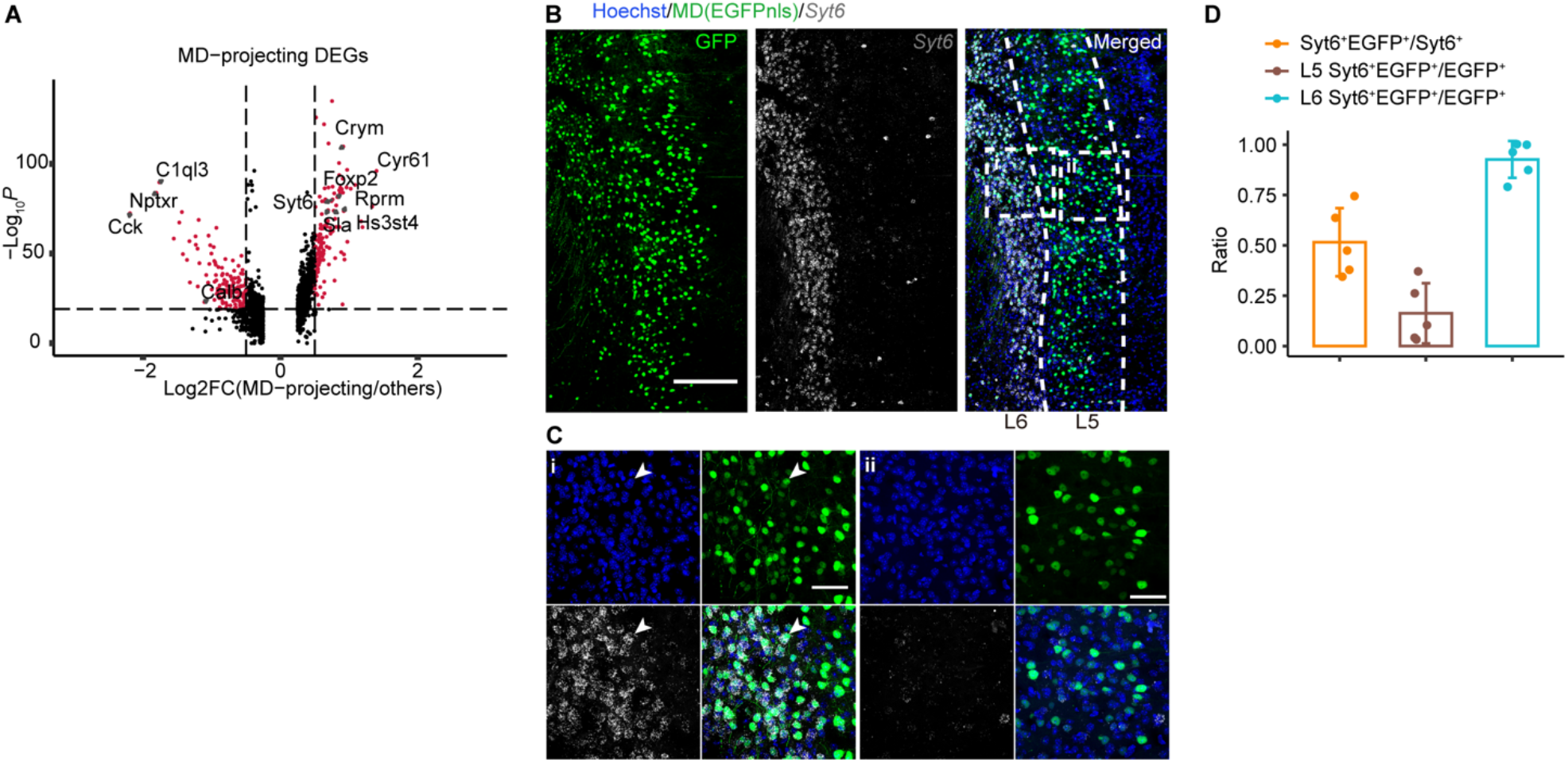
Transcriptional profiling of projection target-specific vmPFC neurons. (**A**) Volcano plots DEGs of MD-projecting versus non-MD-projecting vmPFC neurons. Assigned DEGs (red dots) were determined using threshold: Log_2_ fold change=0.5, p value cutoff= 10^−20^. (**B**) Immunostaining of EGFP (MD) and tdTomato (LH), and RNA FISH of *Syt6*. (**C**) Enlarged view of dotted box in (**B**). (i) represents typical view at layer 6 and (ii) represents typical view at layer 5. Arrow head indicates *Syt6*^+^EGFP^+^ neurons. (**D**) Quantifications of (**B**). (**B**) Scale bars, 200 μm. (**C**) Scale bars, 50 μm. N = 3 mice. Data are presented as mean ± SD.

We further validated the molecular features of neurons associated with their specific projections by combining RNA fluorescence in situ hybridization (FISH) and retrograde labelling. *Syt6* is one of the DEGs of MD-projecting neurons (***Figure 4B***), and is the marker gene of L6-Syt6 cluster. By retrograde labeling of MD-projecting neurons and *Syt6* FISH experiment, we found that about 56% *Syt6*^+^ neurons project to MD. Further statistical analysis showed that most of the MD-projecting (EGFP^+^) *Syt6*^+^ neurons were located in layer 6 but not layer 5 (***Figure 4F-H***), similar to the pattern obtained in our MERGE-seq analysis (***Figure 2E***). These results are in accordance with single-neuron projectomic and transcriptomic analysis of MERGE-seq, indicating that MERGE-seq can faithfully reveal the transcriptomic features of projecting-specific neurons.

### MERGE-seq uncovers the molecular features of collateral projection neurons in vmPFC

Axons of projection neurons, including vmPFC neurons, have highly complex collaterals, which could regulate information processing and neural response properties at the microcircuit level (Gagnon and Parent, 2014; Gao et al., 2022; Rockland, 2019). However, the molecular features of neurons sending collateral projections remain elusive. MERGE-seq provides an opportunity to explore. Here, we identified DEGs for neurons with dedicated and bifurcated projection pattern (***Figure 5A***). We asked whether there was transcriptional difference between neurons with dedicated projection to A and neurons with bifurcated projection to A and B. DEGs were rare in comparisons between projection patterns A/B vs. A, or A/B vs. B in all of groups we tested, except for the DMS+LH group and DMS+MD group (***Figure 5B, C, Figure5-figure supplement 1***). We found that DMS+LH projection neurons were transcriptionally distinct to DMS but similar to LH, and DMS+MD neurons were transcriptionally distinct to DMS but similar to MD (***Figure 5B, C, Figure5-figure supplement 1***). Specifically, we identified a set of genes which differentially expressed in DMS+LH projection neurons (such as *Pou3f1, Bcl6, Bcl11b*, and *Igfbp4*) or DMS+MD projection neurons (such as *Rprm, Crym, Hs3st4* and *Hsp90ab1*).

**Figure 5.**
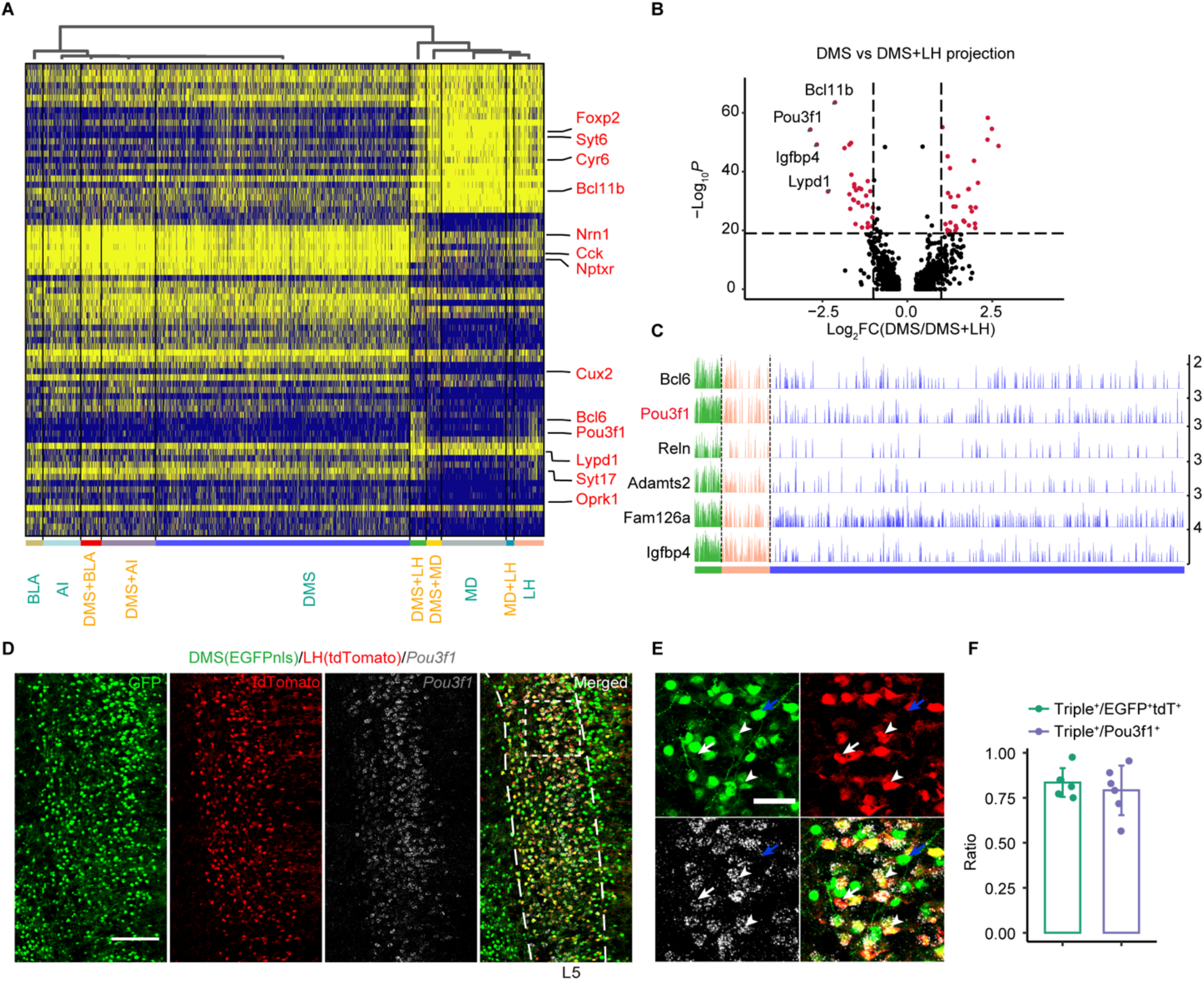
Molecular features of single vmPFC neuron with bifurcated projections to downstream targets. (**A**) Heatmap showing scaled expression of calculated DEGs based on 10 projection patterns. Top 10 DEGs ordered by average log_2_ fold change of each pattern were selected. Typical DEGs are labeled in red. (**B**) Volcano plot showing genes differentially expressed in the DMS/LH group (red spots) compared to the DMS projection pattern. (**C**) Track plots showing normalized data of the selected DEGs in DMS-dedicated, LH-dedicated, and DMS+LH-bifurcated projection pattern. The gene labeled red (*Pou3f1*) was used to perform RNA FISH. (**D**-**F**) Examining *Pou3f1* and DMS+LH projection pattern using RNA FISH and immunostaining of dual-color traced retrograde labeled neurons. Virus injection scheme was the same as in **Fig. 3H**. Scale bars, 200 μm. (**E**) Enlarged view of dotted box in (**D**). Arrow heads indicate *Pou3f1*^+^EGFP^+^tdTomato^+^ neurons, white arrows indicate *Pou3f1*^+^EGFP^-^tdTomato^+^ neurons, and blue arrows indicate *Pou3f1*^-^EGFP^+^tdTomato^-^ neurons. Scale bars, 50 μm. (**F**) Quantification of (**D**). N = 3 mice, Data are presented as mean ± SD.

Interestingly, *Pou3f1* and *Bcl6* are marker genes of L5-Bcl6 neurons (layer 5 neuron subtype), which are the main neuron subtype in DMS+LH projection neuronal population (***Figure 3G***). We next verified the specific gene expression in DMS+LH projection neurons by using RNA FISH in combination with dual-color retrovirus labelling assay (***Figure 5D***). We found that the expression of *Pou3f1* was mainly distributed in layer 5, where *Pou3f1* was specifically expressed in dual-color labeled DMS+LH projecting neurons (white arrowheads, ***Figure 5E***) and LH projecting neurons (white arrows, ***Figure 5E***), but not DMS projecting neurons (blue arrows, ***Figure 5E***). Quantification analysis showed that about 83% DMS+LH-projecting (EGFP^+^tdT^+^) neurons expressed *Pou3f1* and about 79% *Pou3f1*^+^ neurons had collateral projections to DMS and LH (***Figure 5F***). These results are consistent with our prediction based on MERGE-seq data (***Figure 3G***).

Together, by MERGE-seq analysis and experimental validation, we uncovered the specific molecular features of dedicated and bifurcated projection neurons in vmPFC, and demonstrated that *Pou3f1*^+^ neurons in vmPFC have bifurcated projection to both DMS and LH, or dedicated projection to LH.

### Machine learning-based modeling reveals gene clusters for predicting projection patterns

Although many efforts have been made to correlate gene expression with neuronal circuit connectivity (Huang et al., 2020; Sorensen et al., 2015; Sun et al., 2021), the lack of a shared coordinate system for two modalities or limited genes examined reduces the prediction precision. MERGE-seq overcomes these challenges by acquiring high-throughput gene expression and projection pattern in the same neuron (***Figure 6A***). To evaluate potential relationships between the transcriptome and projectome, we used a probabilistic classifier, Naïve Bayes classifier, to predict binary projection patterns for each projection target based on transcription profiles. First, we systematically evaluated the parameters from 2 to 5000 highly variable genes (HVGs) and found less HVGs give better predictions. In fact, we encoded binary projection labels for each projection target (barcoded and unbarcoded) and five set of models (AI, DMS, BLA, LH and MD) were independently trained. Using top 50 HVGs for modeling gave the highest F1 score (harmonic mean between precision and recall), AUC and relatively high predication accuracy (see **Methods, *Figure6-figure supplement* 1A**). Next, we chose top 50 HVGs as features to build the model. As a control model, we chose 50 randomly chosen genes. Five projection targets models were independently trained by splitting cells into training (70%) and test dataset (30%). Using top 50 HVGs also gave rise to significantly better model performance in regarding to prediction accuracy, AUC and F1 score, compared to using random chosen 50 genes (***Figure 6B***). These results suggest that the top 50 HVGs are more informative for predicting and decoding projection patterns.

**Figure 6.**
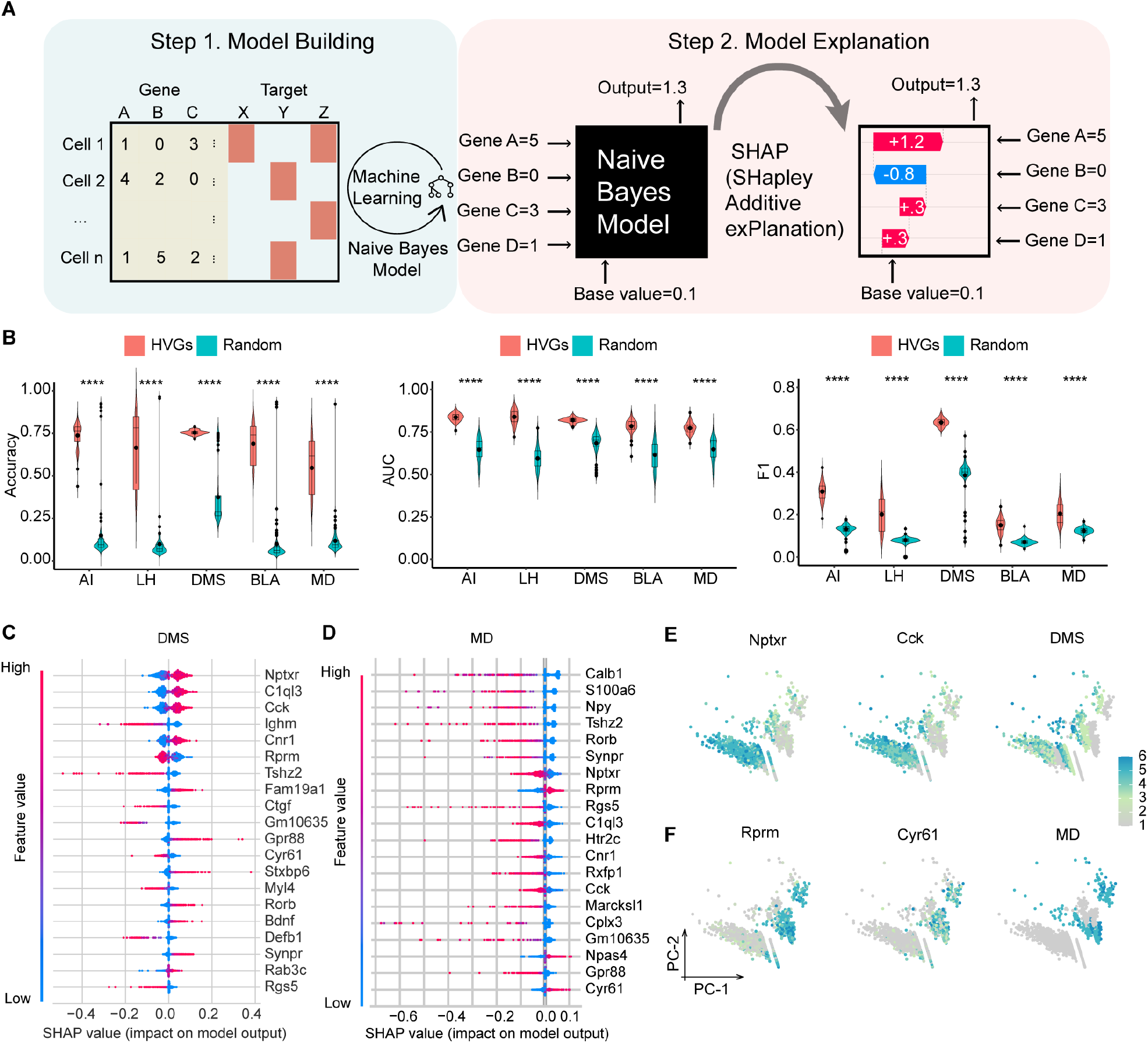
Machine learning-based modeling predicts projection patterns based on gene expression. (**A**) Schematics of machine learning modeling steps. (**B**) Prediction accuracy (left panel), AUC score (middle panel) and F1 score (right panel) of top HVGs and random chosen equal number of genes for modeling building. 100 trials have been performed by randomly sampling 3000 cells from 9368 cells. Top 50 HVGs or 50 randomly chosen genes were used as features per trial. Comparisons were made between models built by the HVGs group and random genes group for each projection target. The displayed P value was computed using a two-sided Wilcoxon test. Data are the mean ± SD. Degree of significance was represented as follows: ****p-value ≤0.0001. (**C, D**) SHAP summary plots of DMS and MD showing important features (genes) with feature effects. For each model, unbarcoded cells were encoded to class 0 and barcoded cells were encoded to class 1. Models were built using top 50 HVGs. (**E, F**) Normalized expression of the most important genes with positive feature effects in Naïve Bayes modeling of DMS (**E**) or MD (**F**) and normalized expression of barcode 1 representing DMS-projecting (**E**) or barcode 4 representing MD-projecting (**F**) on PC1 and PC2 embeddings.

To interpret the important genes contributing to a certain projection pattern, we used a game-theoretic approach to explain the output of HVGs-based Naïve Bayes models (Lundberg et al., 2020) (***Figure 6A***). We used top 50 HVGs to build Naïve Bayes model and summarized effects of HVGs in SHAP (SHapley Additive exPlanations) values for each projection pattern (see **Methods**; ***Figure 6C, D, Figure6-figure supplement 1B-D***). As examples, *Nptxr* and *Cck* genes were the top positive predictors for DMS projection, suggesting that a cell that expresses high levels of *Nptxr* and *Cck* has a higher probability of projecting to DMS. Similarly, *Rprm* and *Cyr61* were the top positive predictors for MD projection. By examining top effective genes (features) on PC embeddings of the projection matrix, we found that the expression pattern of these positive predictors mostly overlaps with projection barcode distribution (***Figure 6E, F***). These results mathematically establish the relationship between gene expression and structural connectivity, indicating the predictive power of a specific gene cluster for projection properties of vmPFC neurons.

## Discussion

Given the complexity of brain circuits, neuronal subtypes must be characterized from multiple viewpoints (Zeng, 2022). Information including neuronal projection patterns (i.e., region-to-region connectivity), physiological properties, gene expression, and how they encode information in behavioral paradigms, are essential to understand functional brain circuits. Therefore, it is inevitably difficult to acquire a complete picture of brain circuits when only one analytic modality is considered. In this study, we have developed a multiplexed barcoding method that is integrated with scRNA-seq, enabling simultaneous transcriptome and projectome analyses. Retrograde AAVs are injected into multiple target regions simultaneously, thereby labelling projection neurons within the brain region of interest and facilitating their transcriptional analysis. Here, by comparing to other methods, we highlight distinct features of MERGE-seq and key biological insights that MERGE-seq can provide.

Early approaches of barcode-based neuronal projection mapping mainly focus on elucidating the projections of individual neurons in a single brain without providing the transcriptional signatures corresponding to those individual neurons (MAPseq, BARseq) (Chen et al., 2019; Kebschull et al., 2016). We therefore developed MERGE-seq to connect single-neuron transcriptome and projectome with high throughput. While there are some conceptual similarities to BARseq or ConnectID (Chen et al., 2019; Klingler et al., 2021), MERGE-seq has its unique features and advantages. BARseq can acquire single-neuron transcriptome and projectome, but with only a number of genes due to limited throughout of in situ sequencing. An improved version of BARseq can allow tens of genes to be detected, but still with a low throughput compared to scRNA-seq and a high cost in regards to synthesizing RNA probes (Sun et al., 2021). ConnectID (scRNA-seq combined with MAPseq) improves the detection of transcriptome using scRNA-seq but has a relatively low recovery rate of cells with transcriptome and projectome simultaneously (∼16%, 391 cells with barcode identity in 2450 cells with scRNA-seq) (Klingler et al., 2021). In contrast, MERGE-seq provides transcriptional profile of thousands of genes in a single-neuron and more than 50% of cells have valid projectome information (barcodes). Another advantage of MERGE-seq is that users only need to sequence one brain region – the source area. While, in BARseq or ConnectID, users need to perform numerous tissue homogenization and sequencing for downstream brain regions and scRNA-seq or in situ sequencing for the source area.

Retro-AAV based-approaches including VECTORseq and Retro-seq were also recently developed to associate neuronal projectome and transcriptome (Cheung et al., 2021; Lui et al., 2021). VECTORseq used several viral transgenes including three recombinase (DreO, Cre, Flpo) and two fluorescent proteins (tdTomato and EGFP) to barcode neurons. However, these transgenes are variable in length (DreO, ∼1000 bp; Cre, ∼ 1000 bp; FLPo, ∼1200 bp; tdTomato, ∼1400 bp; EGFP, ∼700 bp) and driven by different promoters with different strength (EF1a, hSyn, CAG). Such an approach will inevitably result in differential expression of these different transgenes in labelled neurons, which in turn leads to different rates of transgene recovery in these neurons. In addition, viral-mediated overexpression of these recombinase may lead to toxic to the labeled neurons due to non-specific recombination events (Xiao et al., 2012). Therefore, the transgenes used in VECTORseq method should be carefully selected to avoid any potential interferences with neuronal function or gene expression by these different transgenes. In contrast, MERGE-seq used 15-nucleotide barcode sequences in the 3’UTR region of EGFP as projection index driven by the same promoter to label different projection neurons. The expression of these different barcoded EGFP mRNA is comparable, and the number of these barcoded retro-AAV is unlimited. Therefore, MERGE-seq allows users to examine more populations (theoretically unlimited) in one brain and more extensive analysis of collateralization. Further, MERGE-seq can reveal projectome of single collateral projection neurons and identify molecular features of these neurons (***Figure 5***). However, the collateral projection patterns of single neurons were not reported in VECTORseq and Retro-seq-based method (Cheung et al., 2021; Lui et al., 2021).

Lui et al. (Lui et al., 2021) used Retro-seq to investigate the correspondence between transcriptomics and projection patterns of vmPFC neurons. However, Retro-seq used a different strategy than MERGE-seq and provided some similar but mostly distinct biological information. First, Lui et al. (Lui et al., 2021) injected CAV-Cre or retro-AAV-Cre into the specific target region of PFC of Ai14 mice (Rosa-CAG-LSL-tdTomato-WPRE), with each group mice receiving one target region viral injection. Then, they sorted retrogradely labeled tdTomato^+^ neurons from each group mice and performed scRNA-seq separately. In contrast, we injected five distinctly barcoded retro-AAV-EGFP into five different target regions of PFC of the same mouse, with each targeting region receiving one specific barcoded retro-AAV. Second, MERGE-seq but not Retro-seq-based strategy can reveal collateral projection patterns of neurons at single cell resolution. Retro-seq-based strategy inferred collateral projection based on the finding that transcriptome-defined neuron subtypes can project to different targets (or neurons projecting to different targets share common transcriptome-defined neuron subtype). However, the population level multi-target projections of a transcriptome-defined neuron subtype do not necessarily reflect collateral projection of individual neurons within a subtype. For instance, individual neurons within a subtype could project to distinct targets (dedicated projection), but their collective projections show multiple targets. In contrast, in MERGE-seq, individual neurons that were retrogradely labeled multiple barcodes are determined as collateral projection neurons. By MERGE-seq analysis, we uncovered dedicated and collateral projection patterns of individual vmPFC neurons to the five downstream targets, and revealed molecular features associated with these dedicated or collateral projection neurons (***Figure 3-5***). Third, MERGE-seq strategy can be readily applied to other animal models. The requirement of transgenic animals may limit the application of Retro-seq in other animal model where genetic manipulation is difficult. In contrast, in MERGE-seq strategy, we only need to sequence one brain region. This strategy can be readily applied to other animal models such as nonhuman primates.

Of note, despite of chosen different PFC target brain regions, Lui et al. (Lui et al., 2021) and we also provided some similar biological information. Both two studies found that, at the population level, most targets received projections from multiple transcriptome-defined neuron subtypes in PFC, and most transcriptome-defined neuron subtypes in PFC projected to multiple targets, and different transcriptome-defined neuron subtypes had different biased projection pattern. By MERGE-seq analysis, we further extended these findings by showing that most of the dedicated or collateral projections were also from multiple transcriptome-defined neuron subtypes in PFC, and individual neurons in the transcriptome-defined cell subtypes in PFC could have dedicated or collateral projections. These results highlight the complex divergence and convergence of projection in PFC circuits.

A potential technical concern inherited from retro-AAV is that retrograde labelling efficiency of neurons in the target region and the recovery rate of barcoded neuron (single-cell capture or projection index barcode recovery) is inevitably not 100%, which may underestimate or overestimate the projection patterns and cell composition diversity inferred. However, the overall dedicated and collateral projection pattern (within the five targets we examined) and their associated transcriptome will not be greatly affected by the labelling efficiency or recovery rate, and we have experimentally verified some of those collateral projection and their molecular features inferred in our MERGE-seq analysis. Furthermore, carefully controlling the viral titer and refining the procedures of single neuron suspension preparation, as performed in this study, is required to control the labelling efficiency and recovery rate. Notably, the five injection sites we selected are spatially separated, including cortical regions and subcortical regions, and ranging from anterior (Bregma, +2 mm) to posterior (Bregma, -1.5 mm). However, when it comes to study spatially close brain nucleus as downstream targets, for example, several small nuclei in hypothalamus, it will require extensive quantification of virus injection volume to avoid leakage of virus to nearby brain nucleus that are also to be investigated.

In summary, we develop MERGE-seq, a powerful multiplexed projectome and transcriptome analysis platform that will help researchers perform big-data research at low cost. This will enable researchers to understand organizing principles and molecular features of neural circuits across modalities, and to construct more comprehensive mesoscale connectomes.

## Materials and Methods

### AAV vector design

Plasmid pAAV-CAG-tdTomato (Addgene, #59462) was first modified by replacing tdTomato and WPRE with EGFP by T4 DNA Ligase mediated ligation. A 15-bp barcode sequence was then inserted after the stop codon of EGFP, linked by EcoRI restriction enzyme recognition site. Sequences barcode 0 representing the AI target, *CTGCACCGACGCATT*; barcode 1 (DMS target), *GAAGGCACAGACTTT*; barcode 2 (MD target), *GTTGGCTGCAATCCA*; barcode 3 (BLA target), *AAGACGCCGTCGCAA*; barcode 4 (LH target), *TATTCGGAGGACGAC*. Other barcode sequences used for IHC include barcode 10, *AGCTATGCACGATCA*; barcode 206, *GCGTAAGTCTCCTTG*; barcode 210, *CCTGTATGCGTGGAG*. Engineered viruses were produced by Gene Editing Core Facility, Center for Excellence in Brain Science and Intelligence Technology.

### Virus injection

Male adult C57BL/6 mice (8 weeks of age) were anesthetized intraperitoneally using pentobarbital sodium (10 mg/mL, 120 mg/kg b.w.) and unilaterally injected with rAAV2-retro-EGFP-Barcode virus (barcode 0, 1, 2, 3, 4) into five projection targets simultaneously. Coordinates for these injections are as follows. Reference from Bregma and dura, AI at two locations (in mm: 2.0 AP, 2.52 ML, -2.0 DV; 1.6 AP, 2.97 ML, -2.2 DV) with rAAV2-retro-EGFP-barcode 0 (250 nl and 200 nl, 2.90 × 10^13^ VG/ml); DMS at one location (in mm: 0.6 AP, 1.8 ML, -2.2 DV, 8-degree angle), with rAAV2-retro-EGFP-barcode 1 (500 nl, 1.00 × 10^13^ VG/ml); MD at one location (in mm: -1.25 AP, 1.35 ML, -3.55 DV, 20-degree angle), with rAAV2-retro-EGFP-barcode 2 (300 nl, 1.27 × 10^13^ VG/ml); BLA at one location (in mm: -1.5 AP, 3.2 ML, -4.2 DV), with rAAV2-retro-EGFP-barcode 3 (300 nl, 2.00 × 10^13^ VG/ml); LH at one location (in mm: -0.94 AP, 1.2 ML, -4.55 DV), with rAAV2-retro-EGFP-barcode 4 (250 nl, 2.25 × 10^13^ VG/ml). Following each injection, the micropipette was left in the tissue for 10 min before being slowly withdrawn to prevent virus spilling and backflow. Mice were sacrificed 6 weeks after virus injection. Single-cell suspensions were generated as described in methods below.

For dual-color retrograde virus tracing, two locations were unilaterally injected with virus at the same time, one with rAAV2-retro-EGFP-barcode 10 (2.00 × 10^13^ VG/ml) or rAAV2-retro-EGFPnls-barcode 206 or 210 (3.10 × 10^13^ VG/ml for barcode 206 and 4.38 × 10^13^ VG/ml for barcode 210) and one with rAAV2-retro-tdTomato (2.25 × 10^13^ VG/ml). rAAV2-retro-EGFPnls was used to avoid dense fiber staining when performing immunohistochemistry. We deposited the virus plasmid constructs to Addgene (pAAV-CAG-EGFP barcode-(0-10)-SV40 polyA, pAAV-CAG-EGFPnls barcode-(206, 210)-SV40 polyA; Addgene ID 190864-190876).

### scRNA-seq sample and library preparation

For mice without FAC-sorting (mouse 1, 2, 3), three mice that had been injected with virus were anaesthetized and then subjected to transcranial perfusion with ice-cold oxygenated self-made dissection buffer (in mM: 92 Choline chloride, 2.5 KCl, 1.2 NaH_2_PO_4_, 30 NaHCO_3_, 20 HEPES, 25 Glucose, 5 Sodium ascorbate, 2 Thiourea, 3 Sodium pyruvate, 10 MgSO_4_.7H_2_O, 0.5 CaCl_2_.2H_2_O, 12 N-Acetyl-L-Cysteine). The brain was removed, 300 μm vibratome sections were collected, and the PrL and IL regions were microdissected under a stereo microscope with a cooled platform. Brain slices were incubated in dissection buffer with 10 μM AMPA receptor antagonist CNQX (Abcam, ab120017) and 50 μM NMDA receptor antagonist D-AP5 (Abcam, ab120003) at 33°C for 30 min. The pieces were dissociated first using the ice-cold oxygenated dissection buffer added papain (20 units/ml, Worthington, LS003126), 0.067 mM 2-mercaptoethanol (Sigma, M6250), 1.1 mM EDTA (Invitrogen, 15575020), L-Cysteine hydrochloride monohydrate (Sigma, C7880) and Deoxyribonuclease I (Sigma, D4527), with 30-40 min enzymatic digestion at 37°C, followed by 30 min 1 mg/ml protease (Sigma, P5147) and 1 mg/ml dispase (Worthington, LS02106) enzymatic digestion at 25°C. Supernatant was removed and digestion was terminated using dissection buffer containing 2% fetal bovine serum (FBS, Bioind, 04-002-1A). Single-cell suspension was generated by manual trituration using fire-polishing Pasteur pipettes and filtered through a 35 μm DM-equilibrated cell strainer (Falcon, 352052). Cells were then pelleted at 400 g for 5 min. The supernatant was carefully removed and resuspended in 1-2 ml dissection buffer containing 2% FBS. The suspension was then subjected to the debris removal step using the Debris Removal Solution (Miltenyi, 130-109-398). Cell pellets were resuspended and 48,000 cells were loaded into 3 lanes to perform 10X Genomics sequencing. For mice with FAC-sorting (mouse 4, 5, 6), PrL and IL regions were microdissected and dissociated as mice without FAC-sorting, cells were sorted to enrich for EGFP-positive rAAV2-retro-EGFP-barcodes labeled cells. About 4893 EGFP-positive cells were captured and loaded to perform 10X Genomics sequencing. Chromium Single Cell 3’ Reagent Kits (v3) were used for library preparation (10X Genomics). Libraries were sequenced on an Illumina Novaseq 6000 system.

### Projection barcode library preparation

Parallel PCR reactions were performed containing 50 ng of post cDNA amplification reaction cleanup material as a template. P5-Read1 (*AATGATACGGCGACCACCGAGATCTACACTCTTTCCCTACACGACGCTC*) and P7-index-Read2-EGFP (*CAAGCAGAAGACGGCATACGAGATAGGATTCGGTGACTGGAGTTCAGACGTGTGCTCTTCCGATCTGgCATGGACGAGCTGTACAAG*) (200 nM each) were used as primers with the NEBNext Ultra II Q5 Master Mix (NEB, M0544L). Amplification was performed using the following PCR protocol: (1) 33°C for 1 min, (2) 98° for 10 s, then 65°C for 75 s (20-24 cycles), (3) 75°C for 5 min. Reactions were re-pooled during 1X SPRI selection (Beckman, B23317), which harvested virus projection barcodes library. 431-437 bp (with 120bp adaptors) libraries were sequenced using Illumina HiSeq X Ten.

### Immunohistochemistry

Mice were sacrificed 6 weeks after virus injection. Mice were transcardially perfused with phosphate-buffered saline (PBS) followed by 4% paraformaldehyde (PFA). Brain samples were extracted and cryoprotected in 20% sucrose/4% PFA, immersed sequentially in 20% sucrose (in 4% PFA) and 30% sucrose (in 0.1 M phosphate buffer, PB) until sunk, and then transferred to 30% sucrose/PB for more than 24 h. Brain samples were flash-frozen on dry ice and sectioned at 30 μm on a cryostat (Leica, SM2010R). For dual-color retrograde virus tracing, brain slices were blocked in 10% donkey serum and 0.3% Triton X-100 at 37°C for 1 h. Slices were then incubated with primary antibodies against green fluorescent protein (GFP, 1:500, Nacalai, 04404-84, RRID: AB_10013361) and tdTomato (1:500, OriGene, AB8181-200, RRID: AB_2722750) at room temperature for 2 h, then 4°C overnight. Slices were washed three times using PBS and incubated with Hoechst 33342 (1:1000, Lifetech, H3570), as well as secondary donkey anti-rat Alexa Fluor 488 antibodies (1:800, Invitrogen, A21208) and donkey anti-goat Alexa Fluor 568 antibodies (1:800, Invitrogen, A11057) at room temperature for 1 h. Slices were washed three times using PBS and coverslipped. Stained slices were imaged with a 4X objective with numerical aperture 0.16 as a map, followed by 1.5 μm increment z stacks with a 10X objective with numerical aperture 0.4 (FV3000, OLYMPUS). Composite images were automatically stitched in the X-Y plane using ImageJ/FIJI. RNA FISH experiments were performed using RNA-Scope reagents and protocols (ACD Bioscience, CA), following instructions for fixed-frozen tissue. For experiments using RNA-Scope, immunohistochemistry was performed following RNA-Scope. Probes of RNA-Scope used in this study include, Mm-Syt6 (449641), Mm-Pou3f1-C2 (436421-C2).

### scRNA-seq data pre-processing

scRNA-seq data were aligned with the modified mouse reference genome mm10-3.0.0 adding five projection barcodes as separate genes. This was only taken as preliminary validation that projection barcodes can be compatible with the 10X Genomics system. Further projection barcode expression was obtained as described in (**Projection barcode library preparation** and **Projection barcode FASTQ alignment**). scRNA-seq data was demultiplexed using the default parameters of Cellranger software (10X Genomics, v3.0.2). Obtained filtered transcription count matrix was used for downstream analysis.

### Projection barcode FASTQ alignment

Demultiplexing of projection index barcode was performed using deMULTIplex R package (v1.0.2) (https://github.com/chris-mcginnis-ucsf/MULTI-seq). Tag parameters in “MULTIseq.preProcess” function were adjusted according to our user-defined position of index barcode length and position.

### scRNA-seq transcriptional expression analysis

The filtered count matrix was analyzed and processed using Seurat and Scanpy, including data filtering, normalization, highly variable genes selection, scaling, dimension reduction, and clustering (*39, 40*). First, scRNA-seq data from 3 samples of unsorted cells and 1 sample of sorted EGFP-positive cells were created as Seurat object separately; genes with less than 3 counts were removed and cells with fewer than 200 genes detected were removed. Second, four Seurat objects were merged using the “merge” function in Seurat. Downstream analysis of merged Seurat objects are as follows: (1) Data filtering: cells with a mitochondrial gene ratio of greater than 20% were excluded. We kept cells for which we detected between 500 and 8000 genes (cells with more than 8000 genes detected were considered potential doublets), and between 1000 and 60000 counts (cells with more than 60000 counts detected were considered potential doublets). (2) Data normalization: for each cell, counts were log normalized with the “NormalizeData” function in Seurat; “scale.factor” was set to 50000. (3) Highly variable gene selection: 2000 highly variable genes were calculated using the “FindVariableFeatures” function in Seurat. (4) Data scaling: the Seurat object was performed using the “ScaleData” function with default parameters. The number of counts, number of genes, mitochondrial gene ratio, and sorting condition were regressed out in “ScaleData”. (5) Principal component analysis: highly variable genes were used to calculate principal components in the “RunPCA” function. 100 principal components (PCs) were obtained and stored in Seurat object for computing neighborhood graphs and uniform manifold approximation and projection (umap) in following section. (6) Leiden clustering: Seurat object was converted into loom file and imported by Scanpy. A neighborhood graph of observations was computed by “scanpy.pp.neighbors” function in Scanpy. Then, leiden algorithm was used to cluster cells by “scanpy.tl.leiden” function in Scanpy. (7) Cluster merge and trimming: The top 200 differentially expressed genes for each cluster were calculated using the “scanpy.tl.rank_genes_groups” function in Scanpy using parameters method=“wilcoxon” and n_genes=200. Cluster annotation was performed manually based on previously reported markers of PFC all cell types, layer, neuron subtypes, and mouse brain atlas (Bhattacherjee et al., 2019; Sorensen et al., 2015). Cell clusters with similar marker genes were merged into one cluster. Complete marker lists for all cell types and all excitatory neuron subtypes calculated using “FindAllMarkers” function in Seurat were provided (see ***Supplementary file 3***).

### Cell type replicability assessment

To evaluate whether the transcriptional cell types we recovered and annotated correlated with previous reports (Bhattacherjee et al., 2019), we applied MetaNeighbor R package (v1.6.0) (*41*) to calculate AUROC score as a performance vector on all cell types and excitatory neuron subtypes. First, we chose 2000 genes as integration features of our Seurat object and public datasets prepared as Seurat object. Then, we integrated two datasets using anchors found by the “FindIntegrationAnchors” function. Next, we prepared a normalized data matrix of 2000 anchor genes (variable genes) as SummarizedExperiment class using the SummarizedExperiment R package (v1.16.1). Fast, low memory, and unsupervised version of MetaNeighbor were used to calculate the AUROC score (MetaNeighborUS function, fast version was implemented). Cross-dataset mean AUROC scores were plotted in the heatmap.

### Binary projection pattern classification

To determine valid barcoded cells, we hypothesized that all the non-neuronal cells (e.g., endothelial cells, oligodendrocytes, astrocytes) would not be transduced by rAAV2-retro (*42*). Next, we calculated empirical cumulative distribution of barcode unique molecular identifier (UMI) by each projection target (AI, DMS, MD, BLA, LH) of non-neuronal cells, i.e., by each distinct barcode (barcode 0, 1, 2, 3, 4). To achieve a stringent categorization of barcoded cells, we calculated a UMI threshold using empirical cumulative distribution function (ECDF) that determines 99.9% of non-neuronal cells as un-barcoded cells. Next, a cell is determined to be validly barcoded if the number of the barcode UMIs within the cell is larger than the threshold.

For example, the calculated threshold of UMIs for barcode 0 (AI) is 23, which means if a cell contains more than 23 UMIs of barcode 0, then this cell is validly barcoded by AI. UMIs threshold for DMS, 71; for MD, 134; for BLA, 9; for LH, 125 It is worth mentioning that the UMI threshold differs for different targets due to different magnitude of barcode expression of each projection target (***Figure 1-figure supplement 1I***). Finally, we dropped UMI counts of determined unbarcoded cells to zero to obtain the index barcode counts matrix used for downstream analysis. Binary projection patterns were calculated by five projection targets set intersections of corresponding barcoded cells. Only the top 10 frequent binary and collateral projection patterns were kept for reliable inference.

### The conditional probability of projection pattern and determination of over- and under-represented projection patterns

In short, we denoted conditional probability *P*(*B*|*A*) as a proportion of neurons projecting to region B (N_AB_) within a subgroup of neurons projecting to region A (N_A_), thus 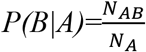. Methods for statistical significance analysis of over- and under-represented projection patterns were described previously (Gergues et al., 2020; Han et al., 2018).

### Projection pattern-specific differentially expressed genes analysis

Differentially expressed genes were calculated using the default parameters of the “FindMarkers” function in Seurat, except the MAST algorithm was used to do DE testing. For the DEG volcano plot, the chosen cut-off for statistical significance was 10^−20^, and chosen cut-off for absolute log_2_ fold-change was 0.5. Volcano plots were implemented using the EnhancedVolcano R package (v1.4.0).

For the DEG heatmap in Figure 5A, the top 10 DEGs ordered by average log_2_ fold-change were chosen from each binary cluster. The heatmap was implemented using the “sc.pl.heatmap” function in Scanpy.

### Machine learning implementation on projection and transcription data

Naïve Bayes was applied to perform a machine learning classification task. We first encoded binary projection labels for each projection target (barcoded and unbarcoded) and five set of models (AI, DMS, BLA, LH and MD) were independently trained. We explored a parameter range of number of the top highly variable genes (HVGs) (2, 5, 10, 20, 50, 100, 200, 300, 400, 500, 1000, 2000, 5000) to fit the model. 3000 cells were randomly sampled from 9368 excitatory neurons and top HVGs were selected by default order of results based on “FindVariableFeatures” function of Seurat per trial. In total, 100 trials were repeated.

To interpret contribution of important genes for each HVGs-based Naïve Bayes model, data matrix for modeling building is constructed as below: for each projection target, 9368 excitatory neurons × (normalized expression of the top 50 HVGs + binary projection labels), or 9368 excitatory neurons× (normalized expression of 50 random genes + binary projection labels). Each data matrix was shuffled first and split by training-testing data in a ratio of 0.7. Machine learning workflow was implanted in pycaret python package (v2.3.4) “pycaret.classification” module. First, for each model, we used “setup” function to initialize the training environment and creates the transformation pipeline by setting “target” parameter to column name of input data matrix corresponding to binary projection labels. Then we used “create_model” function to train and evaluate the performance of a given model by setting “estimator” parameter to “nb” and other parameters by default. We implemented kernel explainer of SHAP python package (v0.40.1) to summarize the effects of genes. SHAP explainer was created using “shap. KernelExplainer(model.predict, training data)” function. SHAP values were calculated using “explainer.shap_values(testing data)” function, and plotted by “shap.summary_plot()” function to create a SHAP beeswarm plot by displaying top 20 features. Training data and testing data for calculating SHAP values were subsampled with 1000 cells.

### Statistical analysis

No statistical methods were used to predetermine sample size. The experiments were not randomized and investigators were not blinded to allocation during experiments and outcome assessment. Two-sided Binomial test was performed and adjusted by Bonferroni correction in Figure 3C, D. Two-sided Wilcoxon test was performed in Figure 6B, Figure 2-figure supplement 1F and Figure 6-figure supplement 1A.

## Data and materials availability

Raw gene expression, barcode count matrices and metadata were available from the Gene Expression Omnibus (GSE210174). The computational code used in the study is available at GitHub (https://github.com/MichaelPeibo/MERGE-seq-analysis) and at Zenodo (https://doi.org/10.5281/zenodo.7189047). The data needed to evaluate the conclusions in the paper can be downloaded at https://figshare.com/projects/High-throughput_mapping_of_single-neuron_projection_and_molecular_features_by_retrograde_barcoded_labelling/150207. All data needed to evaluate the conclusions in the paper are present in the paper and/or the Supplementary Materials.

## Acknowledgments

We thank Dr. Liye Zhang, Pin Wu and Hengxin Liu for generous advice on the bioinformatic analyses.

## Additional information

### Author contributions

J.P., P.X., Z.G.L., and Y.C. designed experiments. J.P. and Z.W. dissected the mouse brains and performed scRNA-seq experiments. T.Y. and H.H. performed the immunohistochemistry experiments. Z.C. performed the retro-barcode virus injection experiments. J.P. and T.L. constructed the retro-AAV plasmids. P.X. analyzed data with considerable input from Z.W. and T.Y.. X.L., L.W., and W.W. helped with bioinformatic analyses. C.T.L., Z.G.L., and Y.C. supervised the project. P.X. and Y.C. wrote and edited the manuscript.

## Supplementary Figures

**Figure 1-Figure supplement 1.**
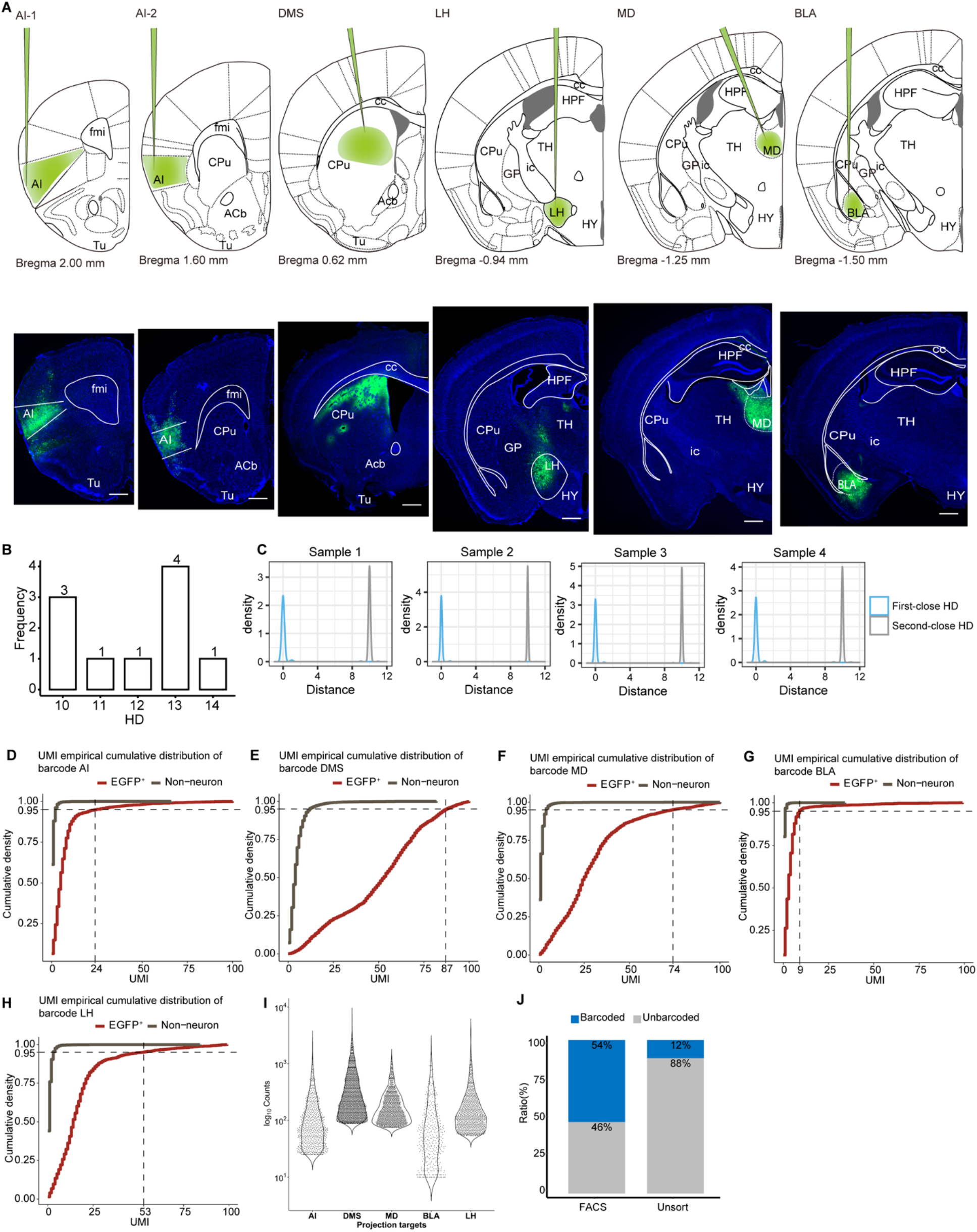
Validation of rAAV2-retro injection sites and determination of valid barcoded cells. (**A**) The position of injection sites (AI-1, AI-2, DMS, LH, MD, BLA) to deliver rAAV2-retro-EGFP plotted on coronal section diagrams (top). The left corner values indicate the anteroposterior distance of the section from bregma. Representative immunohistochemistry images showing rAAV2-retro-EGFP injection sites in coronal sections (bottom). Scale bar: 500 μm. AI-1, agranular insular cortex, anterior; AI-2, agranular insular cortex, posterior. (**B**) Frequency of hamming distance (HD) of five reference barcode sequences. (**C**) Density distribution of HD of best and second-best hit when comparing barcode sequences form reads to and five barcode references. 3 unsorted samples (sample 1-3) and 1 FAC-sorted sample (sample 4). (**D-H**) UMI counts of empirical cumulative distribution for each projection index barcode. Grey lines represent UMI counts of all non-neuron projection index barcode, red lines represent UMI counts of FAC-sorted EGFP^+^ neuron. UMI counts threshold was chosen at the UMI value where the cumulative density of each index barcode UMI counts equaled 0.999 for the non-neuron group (crossing point of horizontal and vertical dashed line). (**I**) Violin plots of Log_10_ normalized projection index barcode counts. Note that after choosing the UMI counts threshold, UMI counts below threshold were dropped to zero. (**J**) Stacked bar plot showing barcoded and unbarcoded cell ratio in FAC-sorted group or unsorted group as determined by stringent UMI threshold.

**Figure 2-Figure supplement 1.**
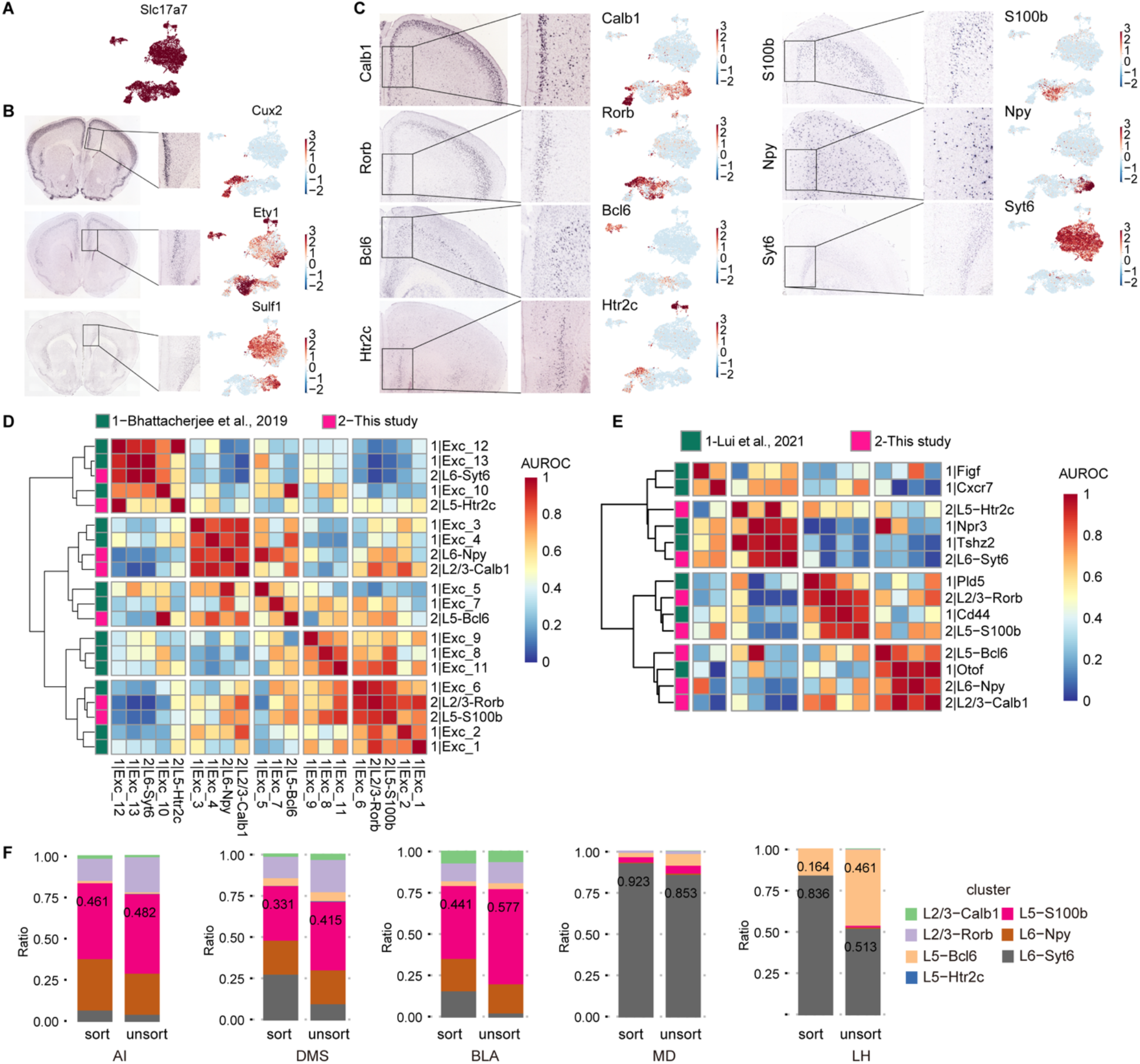
Layer and cluster annotation using the mouse brain atlas and published scRNA-seq transcriptomes, and projection patterns per mouse. (**A**) Normalized *Slc17a7* (*vGlut1*) expression for all extracted excitatory neurons. (**B**) In situ hybridization of typical layer-specific markers within the vmPFC region from the Adult Mouse Brain Atlas. *Cux2* is layer 2/3-specific; *Etv1* is layer 5-specific; *Sulf1* is layer 6-specific (left). Normalized expression of *Cux2, Etv1*, and *Sulf1* at umap embedding (right). (**C**) In situ hybridization of typical neuronal subtype markers in the vmPFC from the Adult Mouse Brain Atlas. *Calb1* and *Rorb* are layer 2/3-specific; *Htr2c* and *S100b* are layer 5-specific; *Bcl6* is around the transition of layer 2/3 and layer 5. *Syt6* and *Npy* are layer 6-specific, though *Npy* is distributed sporadically. Corresponding normalized gene expression embedded in umap is plotted in the right panel. (**D, E**) Heatmap showing correlation between annotated excitatory neuron subtypes recovered in this project and the dataset from Bhattacherjee et al. (Bhattacherjee et al., 2019) and Lui et al. (Lui et al., 2021) (**F**) Stacked bar plots showing neuronal subtype composition of pooled unsorted mouse (unsort 1, 2, 3) and pooled FAC-sorted mice (sort 4, 5, 6) for each projection target. Statistical analysis uses multiple mean comparison by two-sided Wilcoxon test with Holm correction. There is no significant difference between unsort and sort group for each neuronal subtype composition per downstream target.

**Figure 3-Figure supplement 1.**
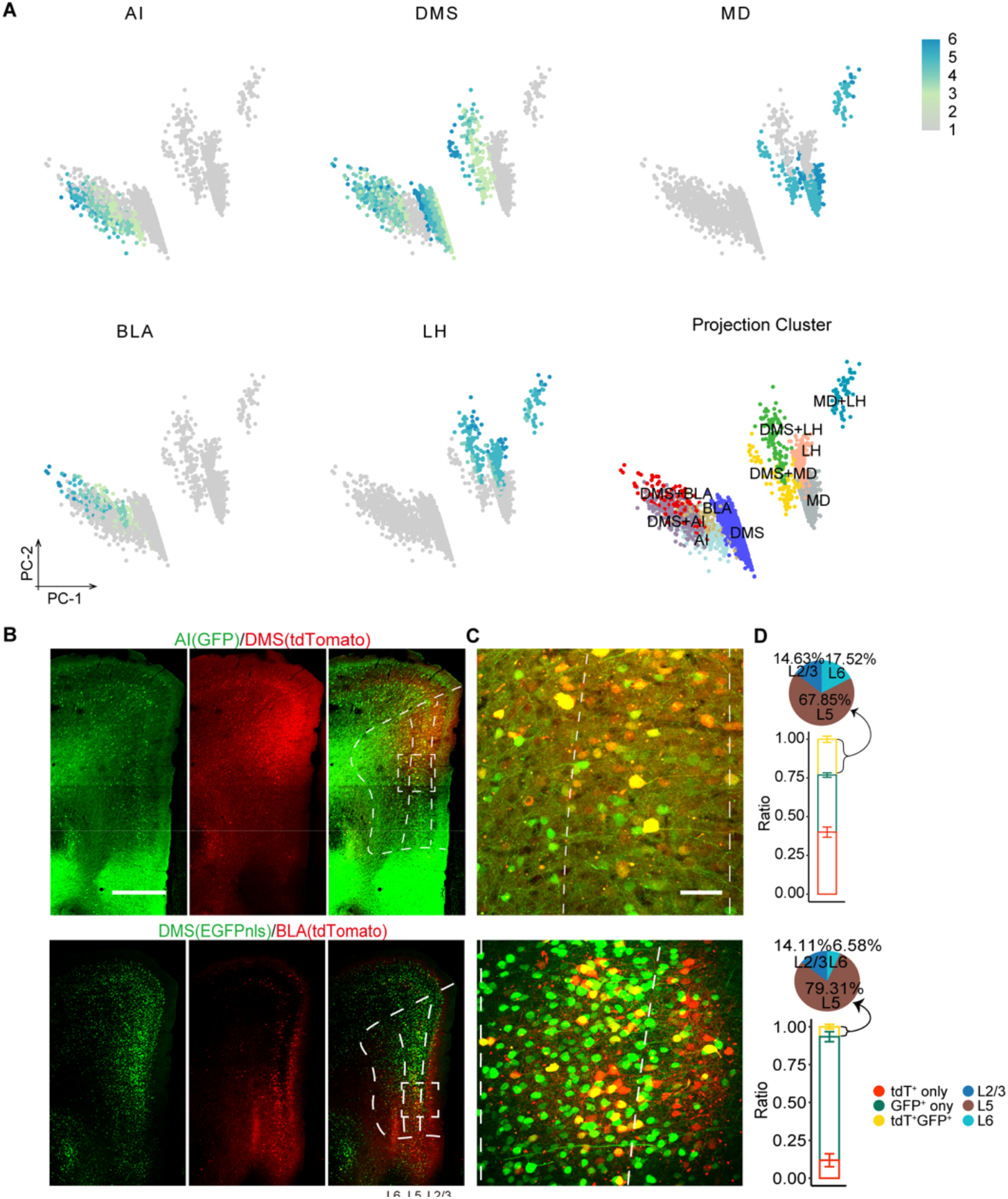
PCA plot of projection clusters, immunostaining of dual-color, retrogradely labeled neurons and quantification. (**A**) Normalized projection index barcode expression on PC1 and PC2 embeddings and binary projection annotation labeled on PC1 and PC2 embeddings. Note that only the 10 most frequent binary projection patterns were included. (**B**) Immunostaining of dual-color retrogradely labeled neurons of AI (GFP) /DMS (tdTomato) and DMS (GFP) /BLA (tdTomato). Dotted line depicts layers 2/3, 5, and 6. Scale bars, 500 μm. (**C**) Enlarged view of dotted box in (**B**). Scale bars, 100 μm. (**D**) Histogram shows quantification for single- (red, green) and double- (yellow) labeled neurons as mean percentages of total retrograde AAV labeled neurons. AI (GFP) /DMS (tdTomato): n=3, DMS (GFP) /BLA (tdTomato): n = 3 mice. Data are the mean ± SD. Co-localization ratio (yellow bar) of each group from the ipsilateral side: AI+DMS = 23.1%±2.03%, DMS+BLA = 6.59%±1.55%. Pie chart showing layer distribution of double- (yellow) labeled neurons.

**Figure 4-Figure supplement 1.**
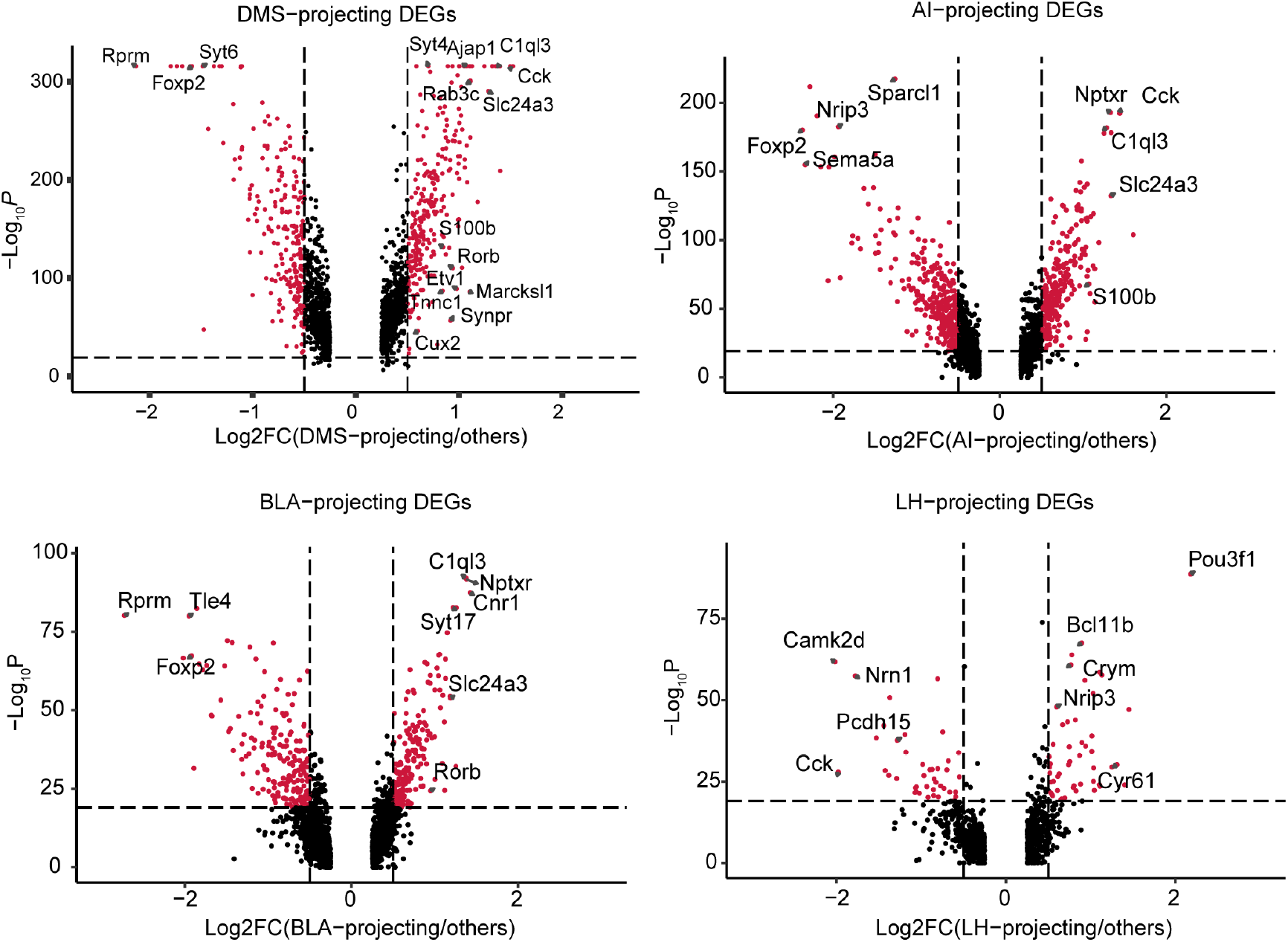
Transcriptional profiling of projection target-specific vmPFC neurons. Volcano plots DEGs of AI-projecting versus non-AI-projecting vmPFC neurons, BLA-projecting versus non-BLA-projecting vmPFC neurons, and LH-projecting versus non-LH-projecting vmPFC neurons. Assigned DEGs (red dots) were determined using threshold: Log2 fold change = 0.5, p value cutoff = 10^−20^.

**Figure 5-Figure supplement 1.**
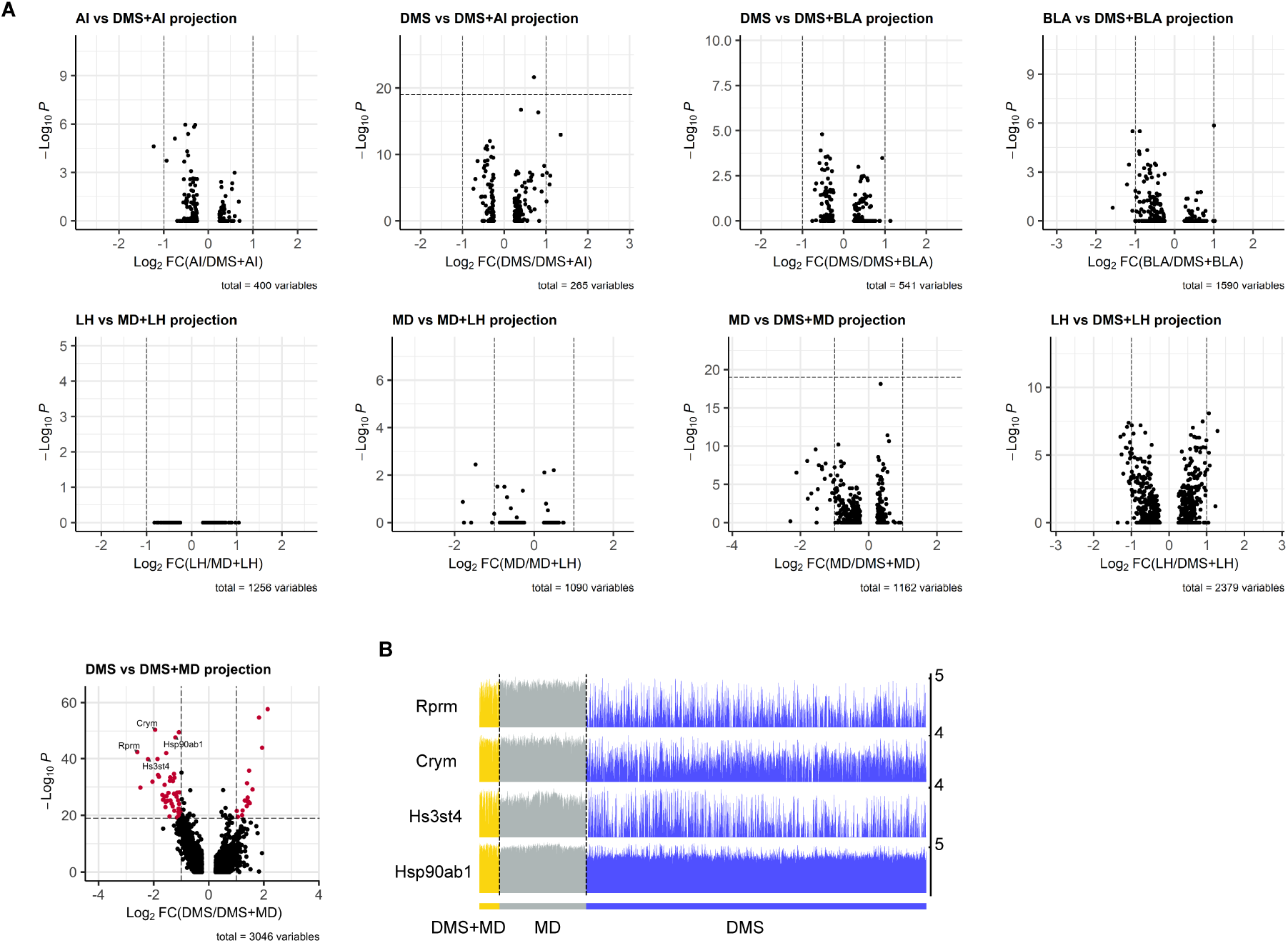
DEGs between dedicated projection neurons versus bifurcated neurons. (**A**) Volcano plots of DEGs calculated between A and A/B projection patterns. See also **Fig. 5**. Assigned DEGs (red dots) were determined using threshold: Log_2_ fold change = 1, p value cutoff = 10^−20^. (**B**) Track plots showing raw counts of the selected DEGs (DMS versus DMS+MD projection) in DMS-dedicated, MD-dedicated, and DMS+MD-bifurcated vmPFC neurons.

**Figure 6-Figure supplement 1.**
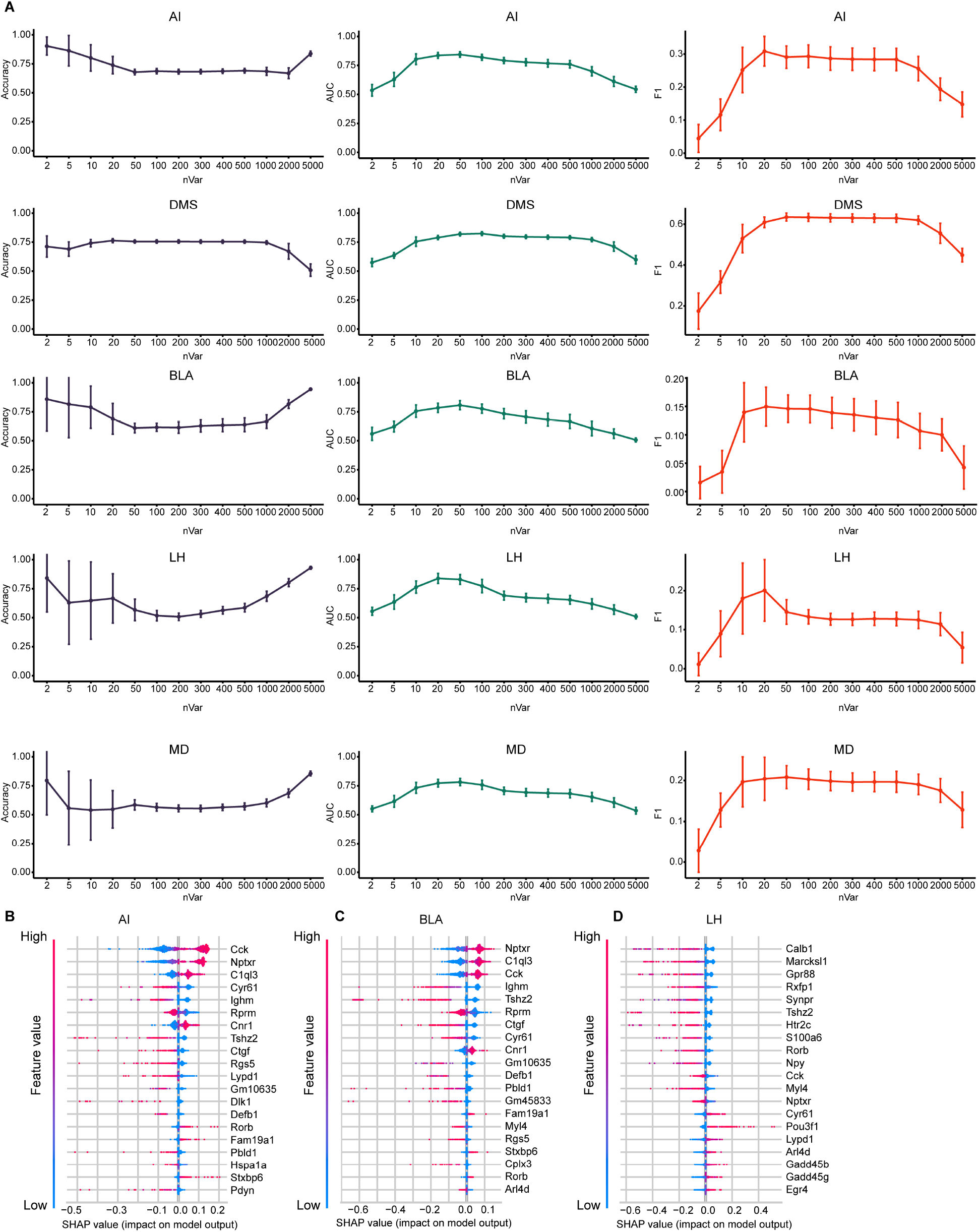
SHAP summary plots of XGBoost models. (**A**) Prediction accuracy (left panel), AUC score (middle panel) and F1 score (right panel) by tuning number of HVGs used for naïve bayes modeling building. 100 trials have been performed by randomly sampling 3000 cells from 9368 cells and calculating top HVGs per trial. (**B-D**) SHAP summary plots of AI, BLA, and LH showing important features (genes) with feature effects. For each model, unbarcoded cells were encoded to class 0 and barcoded cells were encoded to class 1.

